# Transfer learning reveals sequence determinants of the quantitative response to transcription factor dosage

**DOI:** 10.1101/2024.05.28.596078

**Authors:** Sahin Naqvi, Seungsoo Kim, Saman Tabatabaee, Anusri Pampari, Anshul Kundaje, Jonathan K Pritchard, Joanna Wysocka

## Abstract

Deep learning approaches have made significant advances in predicting cell type-specific chromatin patterns from the identity and arrangement of transcription factor (TF) binding motifs. However, most models have been applied in unperturbed contexts, precluding a predictive understanding of how chromatin state responds to TF perturbation. Here, we used transfer learning to train and interpret deep learning models that use DNA sequence to predict, with accuracy approaching experimental reproducibility, how the concentration of two dosage-sensitive TFs (TWIST1, SOX9) affects regulatory element (RE) chromatin accessibility in facial progenitor cells. High-affinity motifs that allow for heterotypic TF co-binding and are concentrated at the center of REs buffer against quantitative changes in TF dosage and strongly predict unperturbed accessibility. In contrast, motifs with low-affinity or homotypic binding distributed throughout REs lead to sensitive responses with minimal contributions to unperturbed accessibility. Both buffering and sensitizing features show signatures of purifying selection. We validated these predictive sequence features using reporter assays and showed that a biophysical model of TF-nucleosome competition can explain the sensitizing effect of low-affinity motifs. Our approach of combining transfer learning and quantitative measurements of the chromatin response to TF dosage therefore represents a powerful method to reveal additional layers of the cis-regulatory code.

## Introduction

Deciphering the cis-regulatory code, the rules by which DNA sequence encodes precise, timely and context-specific gene expression, is a fundamental goal with broad utility in understanding, predicting, and ultimately treating human disease. One key aspect of the cis-regulatory code is chromatin state, the manner in which DNA is packaged in the nucleus. At short length scales (100s to 1000s of base pairs), much of the cellular context-specific chromatin state is determined by transcription factors (TFs), proteins that bind to short DNA sequences termed motifs and, through either mass action or recruitment of enzymes that remodel and modify nucleosomes, set the activity of regulatory elements (REs) that subsequently modulate transcription of target genes^1,2^. Thus, a key part of understanding the cis-regulatory code is the ability to predict chromatin state from the identity and arrangement of TF motifs.

Deep learning models have made substantial progress towards predicting RE chromatin states, as measured by genome-wide assays of histone modifications and chromatin accessibility, from DNA sequence^3–7^. Neural network interpretation tools have revealed that much of this improvement in predictive power derives from flexibly encoding motif affinity and arrangement in a quantitative fashion^6,7^. While such models implicitly learn the concentrations of relevant TFs in a given cell type, the vast majority of models to date have been trained to predict measurements of chromatin state in unperturbed cells, thus precluding a predictive understanding of how chromatin state responds to TF perturbation. Furthermore, TFs have been observed to be highly dosage-sensitive in human variation and disease^8–10^. Using deep learning to understand how chromatin state responds to quantitative changes in TF levels could therefore provide insights into transcriptional regulation while also furthering a mechanistic understanding of how variation in TF levels leads to both normal-range and disease-associated phenotypic diversity.

Predicting the chromatin response to TF dosage presents two main challenges. The first is obtaining relevant experimental measurements for model training. Such measurements involve precise modulation of endogenous TF levels, followed by quantification of chromatin responses. To this end, we recently applied the degradation tag (dTAG) system to achieve tunable modulation of SOX9 dosage in human facial progenitors (cranial neural crest cells, CNCCs) derived *in vitro* from pluripotent stem cells^11^. This revealed a range of chromatin accessibility responses to SOX9 dosage, with most REs being buffered against quantitative SOX9 dosage changes but a subset showing highly sensitive responses. These sensitive responses were selectively linked to specific craniofacial phenotypes associated with SOX9 dosage perturbations, underscoring the importance of dosage effects for understanding human phenotypic variation. The bHLH factor TWIST1 is another compelling candidate for the predictive analysis of dosage effects in CNCCs. Like SOX9, TWIST1 dosage changes are associated with normal-range and disease-related craniofacial variation in humans^12–14^. Moreover, we recently demonstrated that TWIST1 is a key driver of CNCC chromatin accessibility: depletion of TWIST1 in CNCCs results in loss of accessibility at >30,000 REs. In facial progenitors, TWIST1 predominantly binds to a composite motif termed “Coordinator” through cooperative, DNA-guided interactions with homedomain TFs^14^.

The second challenge for predicting the chromatin response to TF dosage is the large number of training examples required by deep learning models. While steady-state models can use all detected REs (∼100s of thousands), only a subset of these REs are expected to respond to perturbation of a single TF (≤10s of thousands), limiting the number of available training examples. Transfer learning, in which deep learning models are “pre-trained” on a larger set of related examples and then fine-tuned to predict the desired task, has recently emerged as an attractive solution to this type of problem, enabling use of deep learning models in data-limited settings^15–19^.

Here, we combined transfer learning of chromatin accessibility models with TF dosage titration by dTAG to learn the sequence logic underlying responsiveness to SOX9 and TWIST1 dosage in CNCCs. Our approach predicted how REs responded to TF dosage, both in terms of magnitude and shape of the response (sensitive or buffered), with accuracy greater than baseline methods and approaching experimental reproducibility. Model interpretation revealed both a TF-shared sequence logic, where composite or discrete motifs allowing for heterotypic TF interactions predict buffered responses, and a TF-specific logic, where low-affinity binding sites for TWIST1 predict sensitive responses. Despite their relatively low importance in models of chromatin accessibility in unperturbed cells, sensitizing sequences show similar levels of conservation to buffering sequences. Finally, we experimentally validated this sequence logic and used biophysical modeling to show that TF-nucleosome competition can explain the sensitizing effects of low-affinity sites.

## Results

### Precise modulation of TWIST1 dosage in hESC-derived CNCCs

We first assessed whether the dTAG system, in which the FKBP12-F36V tag mediates target degradation following addition of the dTAG^V^-1 small molecule, could be used to precisely modulate TWIST1 dosage in human embryonic stem cell (hESC)-derived CNCCs, as we previously did for SOX9^11^. We used a previously-generated hESC line with biallelic knock-in of a FKBP12-F36V-V5 tag at the *TWIST1* N terminus^14^. We differentiated *TWIST1*-tagged hESCs using an established protocol^20,21^ and subsequently titrated *TWIST1* levels by adding varying dTAG^V^-1 concentrations (Figure 1A). As *TWIST1* was not tagged with a fluorescent protein, we measured TWIST1 protein levels by intracellular staining with a monoclonal V5 antibody followed by flow cytometry. We confirmed linearity between intracellular V5 staining intensity and protein abundance by analyzing *SOX9*-tagged CNCCs, which also have the fluorescent mNeonGreen tag (Figure S1A). We achieved five distinct TWIST1 dosages after 24h dTAG^V^-1 treatment, with unimodal single-cell distributions that shifted uniformly with increasing dTAG^V^-1 concentration (Figure 1B, S1B).

**Figure 1.**
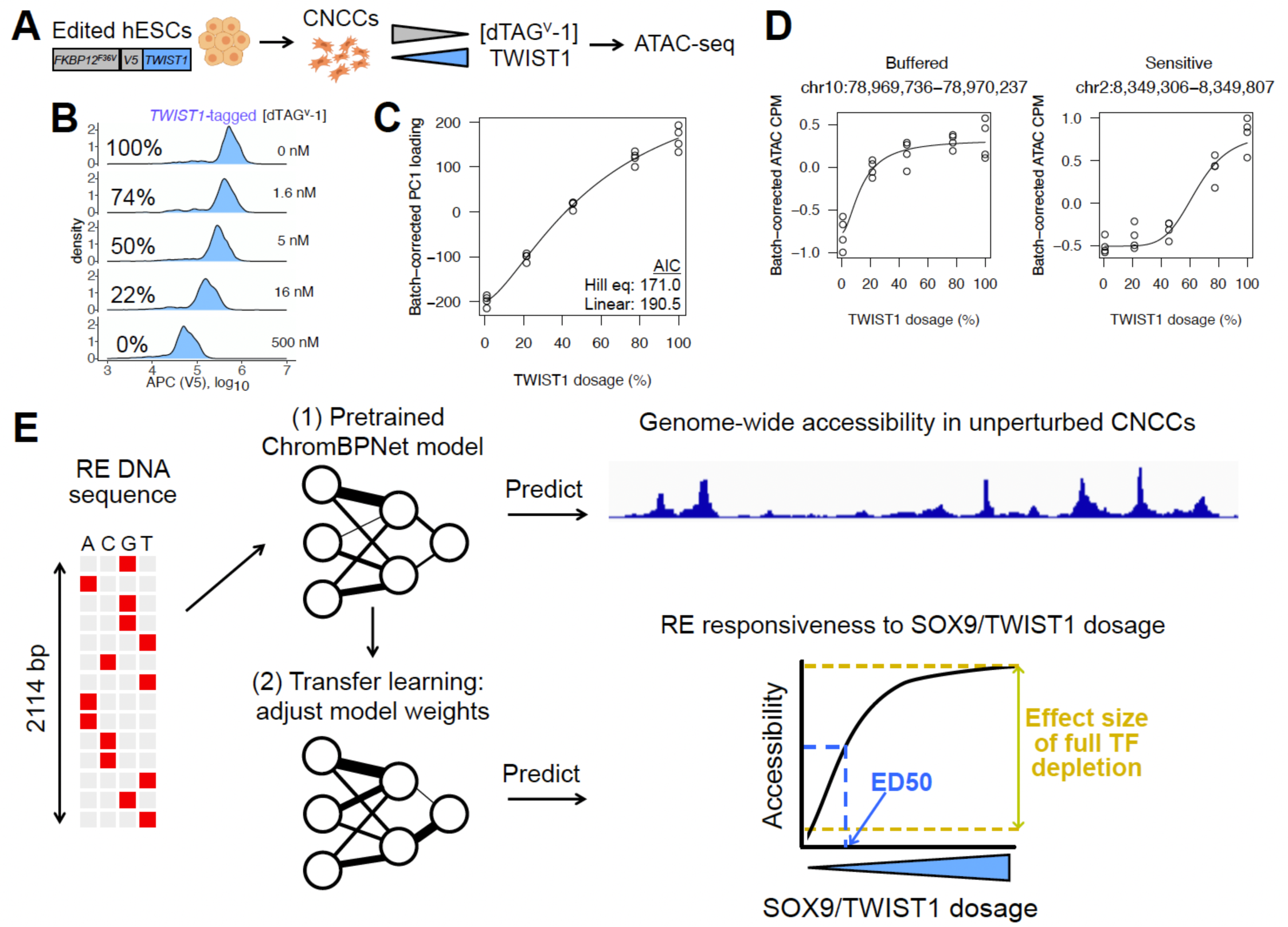
Approach to quantify and predict RE response to TF dosage. (A) Schematic of approach for precise modulation of TWIST1 dosage. (B) Flow cytometry analysis of V5 staining intensity at 24h in *TWIST1*-tagged CNCCs as a function of increasing dTAG^V^-1 concentrations, representative of two independent experiments. (C) Loadings from PC analysis of ATAC-seq CPM of all 151,457 REs across all CNCC samples corrected for differentiation batch and plotted as a function of estimated relative TWIST1 dosage (shown as percentage relative to no dTAG^V^-1). (D) Examples of buffered and sensitive responses, with fitted Hill equation plotted. (E) Schematic of transfer learning approach to predict effect size of full depletion or ED50 for RE chromatin accessibility in response to SOX9 or TWIST1 dosage.

To assess the effect of TWIST1 dosage changes on chromatin accessibility, we carried out the assay for transposase-accessible chromatin with sequencing (ATAC-seq) on *TWIST1*-tagged CNCCs with five different TWIST1 dosages (four biological replicates at each dosage). We observed a nonlinear, monotonic effect of TWIST1 dosage in principal component space (Figure 1C), and inspection of individual REs revealed distinct responses to TWIST1 dosage ranged from buffered (minimal accessibility changes until TWIST1 dosage is greatly reduced) to sensitive (even small decreases in TWIST1 dosage from 100% leading to corresponding accessibility changes) (Figure 1D). Together, these results indicate that, as previously observed with SOX9, TWIST1 dosage effects on chromatin are largely monotonic, but nonlinear and with shapes that can vary substantially between REs.

### Accurate prediction of RE responsiveness to TF dosage by transfer learning

We next sought to use transfer learning to predict RE responsiveness to TWIST1 and SOX9 dosage, using two metrics quantifying the magnitude and shape of the RE response as a proxy for RE responsiveness. First, for all 151,457 ATAC peak regions, we calculated the log_2_ fold-change in accessibility upon full TF depletion. Second, for each RE responding significantly to either SOX9 (35,713 REs as defined in Naqvi et al) or TWIST1 (50,858 REs at 1% FDR) dosage, we calculated the ED50 of a fitted Hill equation. We slightly modified the ED50 calculation from Naqvi et al due to heightened sensitivity of some TWIST1-dependent REs leading to unstable estimates (see Methods) (Table S1,2). Lower ED50 values indicate a more buffered response to decreases in TF dosage from 100%, whereas higher ED50 values indicate a more sensitive response (Figure 1E). While TWIST1-dependent REs typically showed higher ED50 values than SOX9-dependent REs, there was still a substantial variation in both the full depletion effect and ED50 (Figure S2A). Furthermore, among REs dependent on both TWIST1 and SOX9 (18,416 REs; 32,442 and 17,297 REs dependent on only TWIST1 or SOX9, respectively), TWIST1 ED50 values were uncorrelated with SOX9 ED50 values (Figure S2B). Thus, despite extensive co-regulation of REs by both TFs, dosage-sensitivity to individual TFs differs at these shared targets.

We next defined predictive tasks for the full depletion effect and ED50, considering previously observed correlates. For predicting the full depletion effect, we used all 151,457 REs. For predicting ED50, we had previously observed that REs likely directly regulated by SOX9 showed substantially higher ED50 than secondary effects mediated by other downstream TFs^11^. We observed a similar phenomenon for TWIST1 sensitivity, where putative direct targets (defined as TWIST1-bound by ChIP-seq and downregulated upon full depletion), showed higher ED50 values compared to other TWIST1-dependent REs (Figure S2C). We therefore sought to predict ED50 among the directly regulated REs for each TF (9,279 and 21,172 for SOX9 and TWIST1, respectively).

Our transfer learning approach involves pretraining a deep learning model to predict chromatin accessibility levels among ATAC peaks and matched background regions in unperturbed cells, followed by model fine-tuning to predict either the full depletion effect or ED50 for TWIST1 or SOX9 (four prediction tasks) among the above-defined sets of REs (Figure 1E). We used a convolutional neural network (CNN) architecture from the recently developed ChromBPNet model, which provides quantitative predictions and was explicitly designed to account for transposase insertion bias in ATAC-seq^22^. For baseline comparisons, we used regularized (LASSO) linear or random forest regression with standard position weight matrix (PWM) matching of all known motifs as predictors. Because unperturbed chromatin accessibility was correlated with both full depletion effect and ED50 (Figure S2D), we also included this as a predictor in the LASSO and random forest models. We benchmarked all predictions against a lower-bound estimate of experimental reproducibility obtained by comparing full depletion effect or ED50 estimates between two halves of biological replicates.

Fine-tuned ChromBPNet models substantially outperformed baseline approaches and showed prediction accuracy roughly on par with experimental reproducibility (Figure 2A,B). In three of the four prediction tasks (not for SOX9 ED50 prediction), fine-tuned ChromBPNet also outperformed baseline approaches that included real data (chromatin accessibility in unperturbed cells) as a predictor. Omitting either the pretraining or fine-tuning steps resulted in a substantial drop in accuracy (Figure S3A). Prediction accuracy was stable across replicate training runs and independent training-validation-testing data splits (Figure S3B). While the absolute predictive accuracy of the full depletion effects was ∼55-60% higher than that of ED50, this is driven by two technical factors. First, the full depletion predictions are over all 151,457 REs, whereas ED50 predictions are for the much smaller subset of likely direct targets of SOX9 and TWIST1. Indeed, when subsetting the full depletion effect predictions to these likely direct targets, the accuracy improvement over ED50 predictions is lower (∼15-43%, Figure S3C). Second, the full depletion estimate is itself less noisy than the ED50 estimate, as indicated by the higher correlation between replicate splits (Figure 2A,B). Nonetheless, these combined results indicate that our transfer learning approach can accurately predict both the magnitude and shape of the quantitative response to TF dosage.

**Figure 2.**
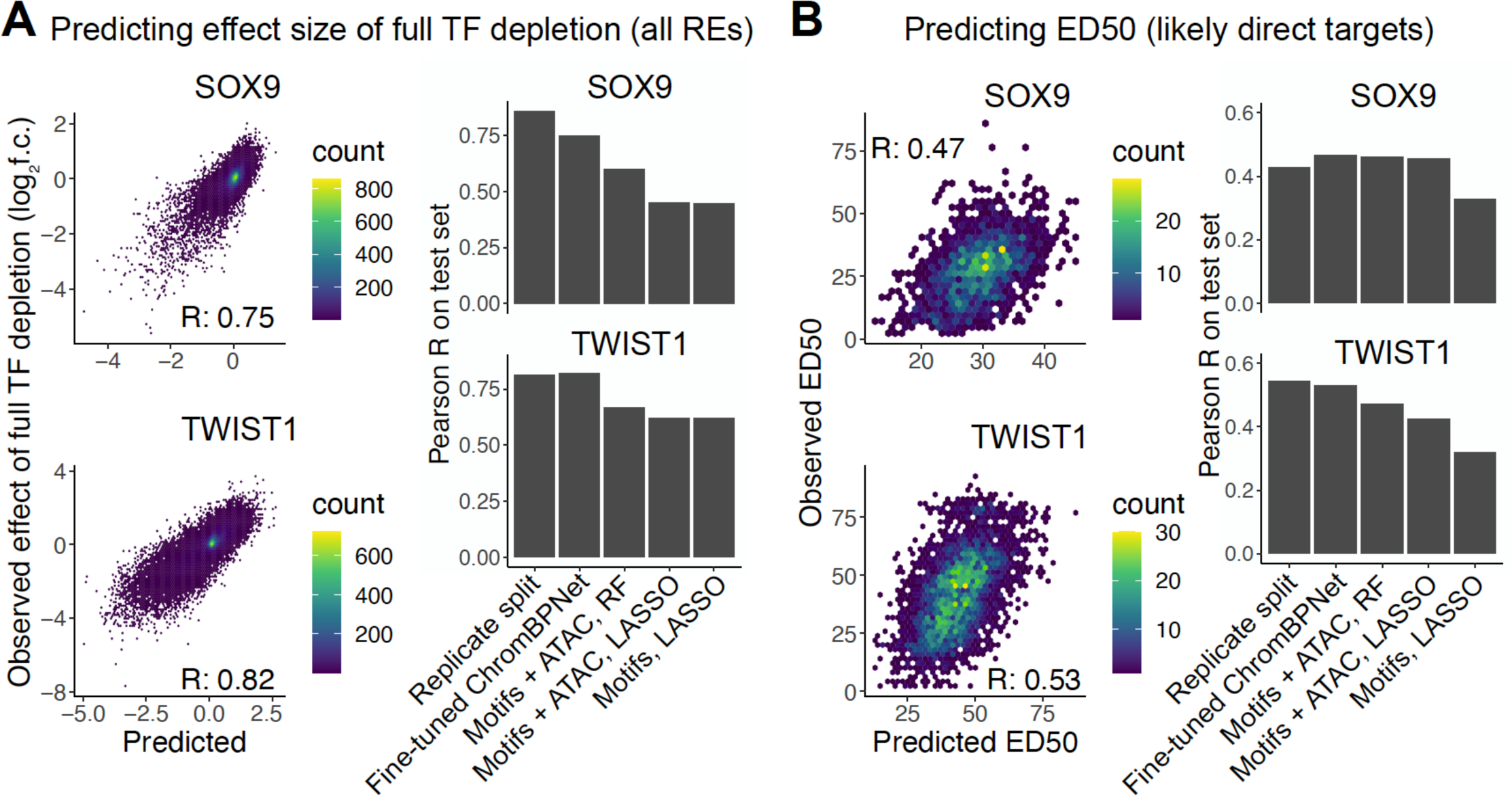
Accurate prediction of the RE response to SOX9 and TWIST1 dosage by transfer learning. (A) Performance of fine-tuned ChromBPNet model (left) on effect of full depletion of SOX9 (top) or TWIST1 (bottom), compared to lower-bound estimate of experimental reproducibility (replicate split), and baseline approaches (right). (B) Same plots as in (A) but for ED50, only considering likely direct targets of SOX9 or TWIST1.

### Sequence features predictive of RE responsiveness to TF dosage

We next sought to interpret the fine-tuned deep learning models to discover the sequence features that underlie their improved performance. We used DeepLIFT^23^ to generate contribution scores, which quantify how each base pair in an RE contributes to the predicted full depletion effect or ED50. We then used Transcription Factor Motif Discovery from Importance Scores (TF-MoDISco)^24^ to discover motifs with high contribution scores, summarized as contribution weight matrices (CWMs). We first assessed sequence features predictive of the full depletion effect, considering all REs active in CNCCs. We and others have previously shown that SOX9 binds a palindrome motif with 3-5 bp spacing, while TWIST1 cooperatively binds the composite Coordinator motif, consisting of a E-box (CANNTG) contacted directly by TWIST1 and homeobox (TAATT[A/G]) sequences separated by a A-rich spacer and bound by a range of homeodomain-containing TFs^11,14,25^. For SOX9, we found that the 3-5bp palindrome motifs all predict a larger loss of accessibility following full depletion (Figure S4A). For TWIST1, both canonical and variant instances of Coordinator predict a larger loss of accessibility (Figure S4B). We expand on these variant motifs further in the analysis of sequence features predictive of ED50. Motifs for specific other TFs (TFAP2, TWIST1, and JUN/FOS for SOX9; TFAP2, SIX, NR2F for TWIST1) predicted gains of accessibility following full depletion, consistent with our previous observations of secondary effects following SOX9 or TWIST1 depletion. E-box motifs specific to the repressive TFs SNAI1/2 (CAGGTG) were also predictive of losses in accessibility, mostly for REs that showed delayed effects following SOX9 depletion (Figure S4A).

We next focused on sequence features predictive of ED50 among the likely direct targets of each TF. For both SOX9 and TWIST1, motifs for TFs other than the perturbed one were predictive of a more buffered response (lower ED50), with the exception of the JUN/FOS motif, which was predictive of sensitivity to SOX9 dosage (Figure 3A,B). In contrast, motifs bound solely by the perturbed TF, such as the SOX9 palindrome and the single or double E-box, were predictive of increased sensitivity (higher ED50) (Figure 3A,B). The observation that buffering is associated with presence of other TF motifs and sensitivity is linked to the homotypic motifs for the perturbed TF recapitulates the patterns we previously observed when analyzing SOX9 ED50 and extends them to be predictive of dosage responses to TWIST1 as well.

**Figure 3.**
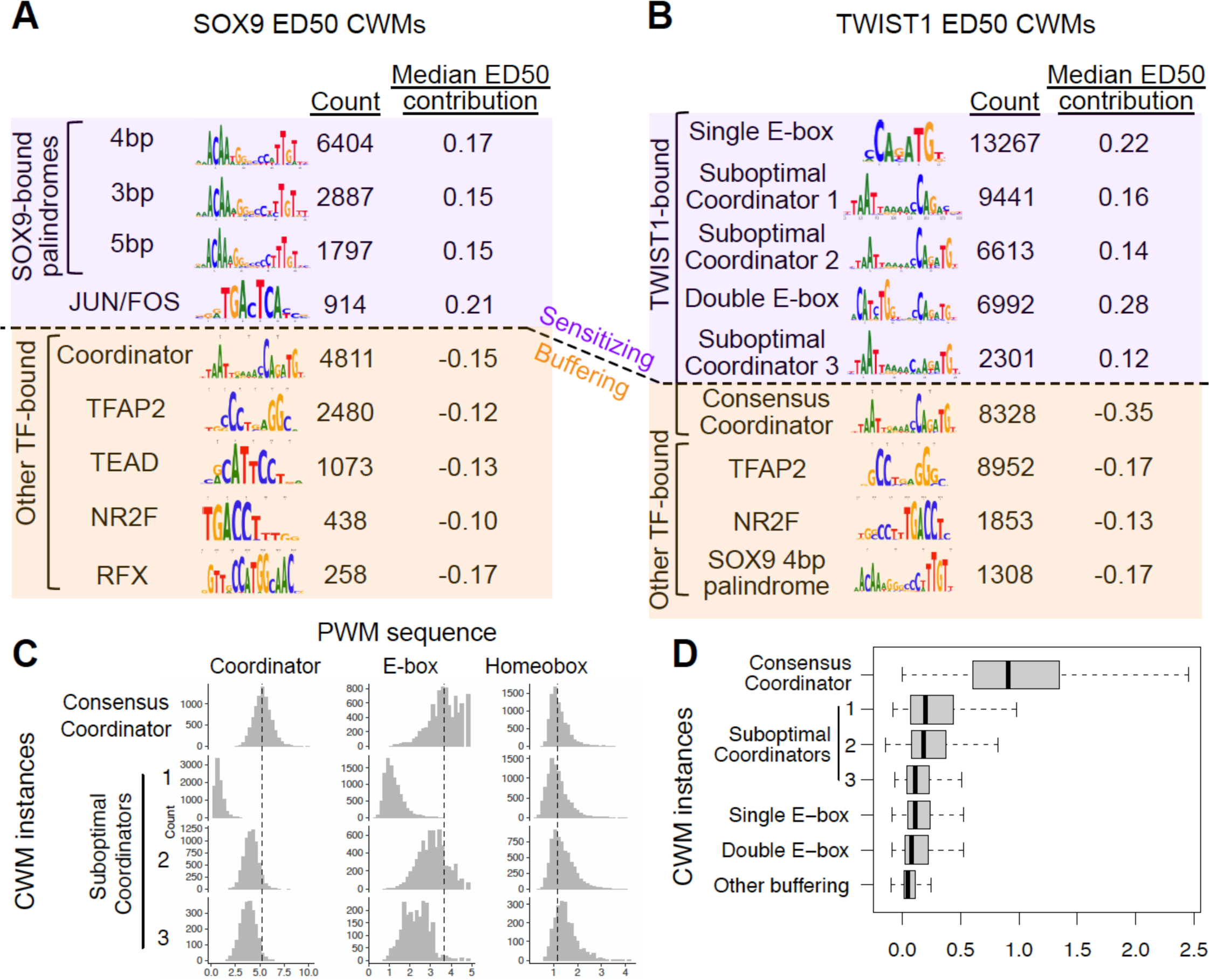
Sequence features predictive of RE sensitivity to SOX9 or TWIST1 dosage. (A,B) Top contribution weight matrices (CWMs) predictive of (A) SOX9 or (B) TWIST1 ED50. Number of individual occurrences of each CWM is indicated under the “count” column, as well as the median of the mean basepair-level contribution scores at all occurrences. “Sensitizing” refers to positive ED50 contributions, whereas “buffering” indicates negative ED50 contributions. (C) For all individual instances of the indicated CWMs predictive of TWIST1 ED50 (rows, i.e. from B), the strength of that CWM’s sequence match to the given position weight matrix (PWM, columns) is shown (x-axis). (D) For the same CWMs, the distribution of contribution to TWIST1 binding, estimated from training a BPNet model on TWIST1 ChIP-seq, is shown.

Surprisingly, however, for TWIST1, diverse types of Coordinator motifs (which are bound by TWIST1 in the E-box portion and homeodomain TFs in the homeobox portion) showed opposing effects on ED50 – CWMs with high similarity to the consensus Coordinator were strongly predictive of buffering, whereas degenerate CWMs lacking preferred nucleotides were predictive of sensitivity. These degenerate CWMs fell into three subclusters: degenerate sequence at the 1) final two or 2) first two base pairs of the Coordinator E-box (CANNNN and NNNNTG), or 3) substitutions throughout the motif (Figure 3B). We have previously found, using electrophoretic mobility shift assays (EMSA), that TWIST1 binds the suboptimal E-box and Coordinator variants containing a single base pair substitution with ∼3-5-fold lower affinity than the canonical Coordinator motif^14^. The Coordinator variants we discovered here have even more degeneracy than the single base pair substitutions previously tested in EMSA, suggesting that even very low-affinity motifs can contribute substantially to TWIST1 responsiveness and confer sensitivity.

To further substantiate the idea that low-affinity binding sites sensitize the response to TWIST1 dosage, we quantified how individual instances of these sensitizing or buffering CWMs match their canonical corresponding PWMs and predict TWIST1 binding. We adapted a previously developed method^6^ to identify individual occurrences of highly contributing sequences and match them to a specific CWM, and then scanned each CWM occurrence with the corresponding PWM model (Table S3,4). Most single or double E-box occurrences sensitizing for TWIST1 showed weak but detectable matches (Figure 3C). The Coordinator CWM instances showed the most stark differences, where most (55.3%) of the sensitizing Coordinator instances are undetectable by even lenient PWM matching thresholds, whereas almost all (92.8%) of buffering Coordinator instances show strong PWM matches (Figure 3C). We then assessed how individual CWM instances contributed to TWIST1 or SOX9 binding, as measured by applying BPNet^7^ to chromatin immunoprecipitation followed by sequencing (ChIP-seq) of each TF. The buffering Coordinator instances were highly predictive of TWIST1 binding, while the degenerate, sensitizing Coordinator instances (as well as the single and double E-boxes) showed less predictive contributions on average but significantly more than other CWMs (Figure 3D). For SOX9, the palindrome sequences showed strong PWM matches and were similar to each other in predicting binding (Figure S4C,D). Together, these results indicate that the chromatin response to TF dosage consists of TF-shared logic, where heterotypic co-binding with other TFs predict buffered responses for both SOX9 and TWIST1, as well as TF-specific logic, where low-affinity TWIST1 binding sites are predictive of sensitivity but high-affinity sites are buffering.

### Buffering and sensitizing motif occurrences show distinct regulatory logic but are similarly constrained

We next assessed distinct and shared features of buffering and sensitizing sequences. There were a total of 22,667 buffering or sensitizing motif occurrences for SOX9 (mean 2.5 per RE) and 62,261 for TWIST1 (mean 3 per RE) across all likely direct target REs. Most REs contained both sensitizing and buffering motif occurrences (Figure 4A,B; Figure S5A,B), indicating that combinations of the two types can concurrently tune RE dosage response. With respect to positioning, sensitizing elements were located further away from the RE summit (point of highest accessibility across the RE) as compared to buffering elements (Figure 4C). We observed similar results when analyzing specific subtypes of sensitizing or buffering motifs (Figure S5C,D). We then compared how these buffering/sensitizing motif occurrences contributed to accessibility levels in unperturbed cells as estimated from the pretrained ChromBPNet model. Inspecting contribution tracks at individual loci revealed many sensitizing instances that would not be detected in the pretrained model of unperturbed accessibility, in contrast to buffering instances which tracked with the unperturbed accessibility contributions (Figure 4D). Indeed, for both SOX9 and TWIST1, buffering sequences showed strong, positive contributions to unperturbed accessibility, while sensitizing sequences showed weaker contributions (Figure 4E).

**Figure 4.**
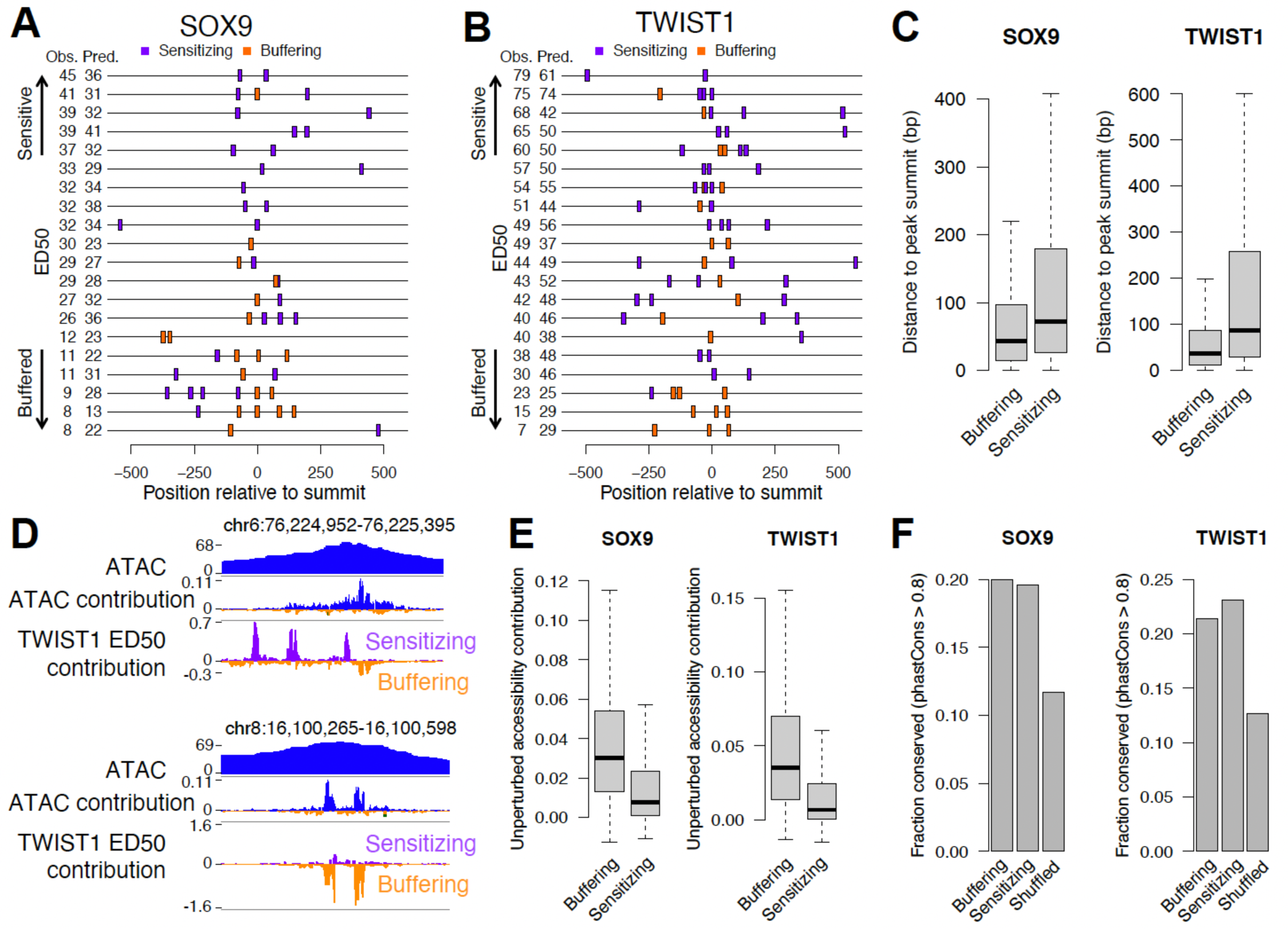
Distinct and shared features of sensitizing and buffering sequences. (A,B) 15 randomly sampled SOX9 (A) or TWIST1 (B) target REs with buffering sensitizing CWM occurrences (colors) indicated. (C) Distance to ATAC peak summit of all sensitizing or buffering CWM occurrences for SOX9 (left) or TWIST1 (right). (D) Examples of sensitizing and buffering occurrences for TWIST1. (E) Accessibility contributions in unperturbed cells of all sensitizing or buffering CWM occurrences for SOX9 (left) or TWIST1 (right). (F) Fraction of buffering, sensitizing, or location-shuffled occurrences showing evidence of evolutionary conservation for SOX9 (left) or TWIST1 (right). Comparisons in C,E p < 2.2e-16 by Wilcoxon rank-sum test. N for groups in C,E,F: SOX9 buffering 9,953; SOX9 sensitizing 12,716; SOX9 shuffled 26,596; TWIST1 buffering 23,647; TWIST1 sensitizing 38,614; TWIST1 shuffled 71,012.

The weaker unperturbed accessibility contributions and distributed positioning of sensitizing elements raises the question of whether they are as biologically relevant as buffering elements. To assess this, we compared signatures of selection between buffering and sensitizing elements, using shuffled regions within the same REs as a control. We considered base pair-level estimates of negative selection from multiple vertebrate genome alignments, as well as joint estimates combining between-species divergence with human polymorphism data^26^. Sensitizing and buffering elements showed very similar degrees of negative selection and both were higher than shuffled regions (Figure 4F, Figure S6A,B). Neither class of element showed evidence for positive selection (Figure S6C). These results suggest that sensitive TF dosage responses, as well as the low-affinity and homotypic motifs that predict them, contribute to organismal fitness and are thus evolutionarily constrained.

### Enhancer reporter assays validate model predictions on mutant sequences

We sought to experimentally validate the sensitizing and buffering sequence elements revealed by the fine-tuned ChromBPNet model. We used enhancer reporter assays, in which an RE is cloned upstream of a minimal promoter and a luciferase reporter gene and assayed for enhancer activity in CNCCs with distinct TWIST1 or SOX9 dosage achieved using dTAG (Figure 5A). While there are known differences between such assays that test ability to activate reporter transcription and the endogenous measures of chromatin accessibility we used to fine-tune ChromBPNet models, previous studies have found RE accessibility to be among the most predictive biochemical features for reporter activity^27^, and the two classes of assays share broadly similar predictive sequence features^15^. We first assayed wild type sequences of 19 REs (12 TWIST1-dependent, 7 SOX9-dependent), observing a positive correlation between endogenous RE accessibility and reporter activity at 100% TF dosage (Figure S7A, Table S5). For the 15 REs that showed significant enhancer activity at 100% TF dosage, we observed a positive correlation between ED50 of endogenous accessibility and the same metrics for enhancer reporter activity (Figure 5B). We observed a similar correlation for the full depletion effect (Figure S7B). These results indicate that episomal reporter assays can recapitulate differences in TF responsiveness observed from endogenous chromatin accessibility measurements.

**Figure 5.**
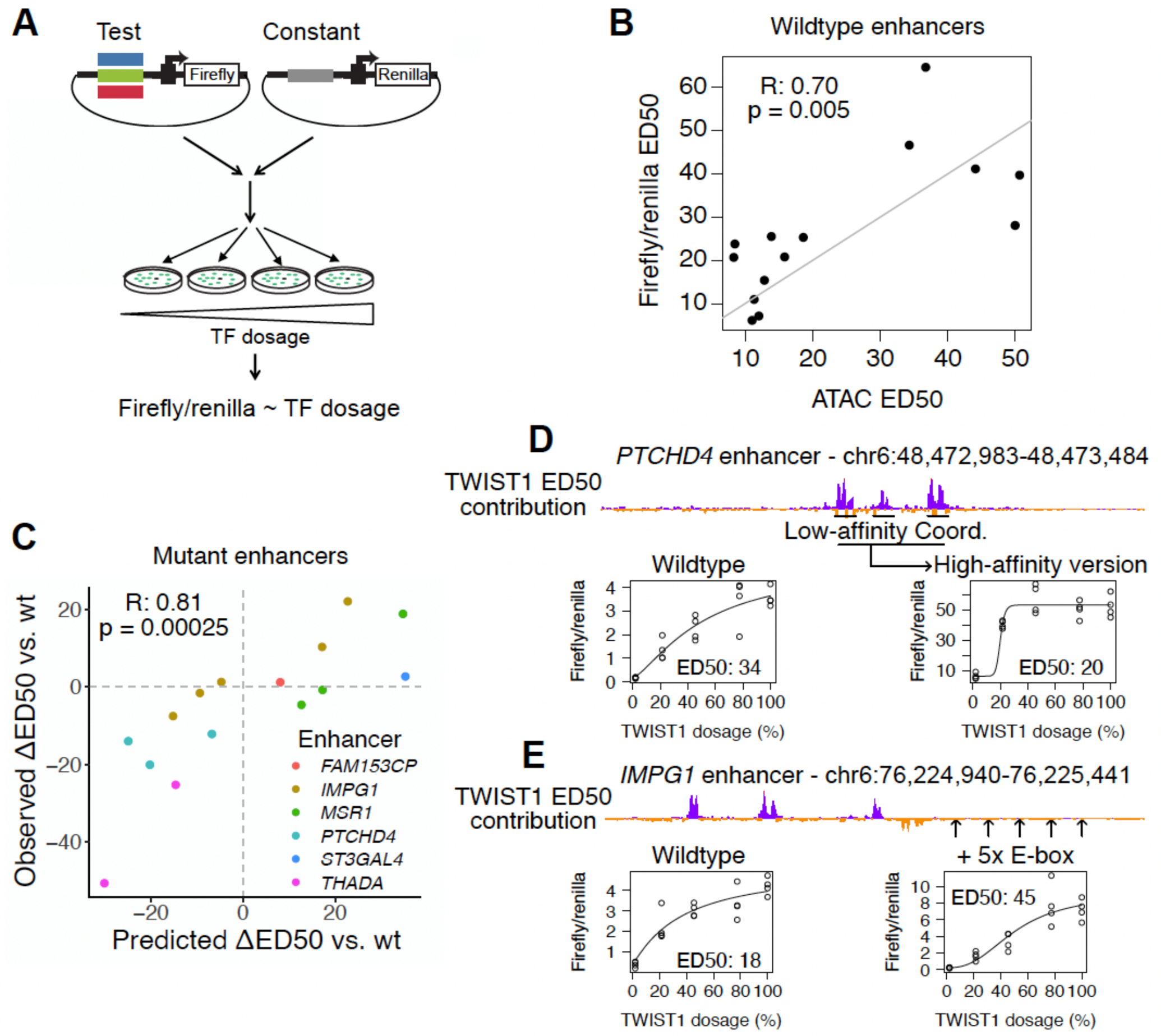
Experimental validation of model predictions by enhancer reporter assays. (A) Schematic of enhancer reporter assay approach. (B) Comparison of ED50 with respect to SOX9 or TWIST1 dosage measured endogenously with ATAC-seq (x-axis) and assessed in the episomal reporter assay (y-axis). Each point is a different wildtype enhancer sequence, ED50 was assessed with at least 4 replicates. (C) Predicted (x-axis) versus observed (y-axis) effect of model-guided mutations on ED50 of various enhancers (colors). ED50 was assessed with at least 4 replicates. (D) Example of *PTCHD4* enhancer, where converting three low-affinity Coordinators/single E-box (peaks in the ED50 contribution track) to high-affinity Coordinators has a buffering effect. (E) Example of *IMPG1* enhancer, where ectopically inserting five single E-box motifs at the indicated positions has a sensitizing effect.

We next tested the effect of sensitizing and buffering features by assaying mutant enhancer sequences predicted to increase or decrease ED50. We selected 7 wild type enhancers with a range of ED50 values and designed mutant sequences, guided by principles from our model interpretation. We designed minimal substitutions to modulate TWIST1 binding site affinity and composition, as well as larger changes involving scrambling or insertion of sensitizing elements at various positions (Table S6). We designed mutant sequences to retain accessibility/enhancer activity such that their ED50 could still be assayed, resulting in 15 mutant sequences from the 7 wild type enhancers. We tested each mutant in the above-described enhancer reporter assay in parallel with its corresponding wild type sequence, calculating the difference in ED50 between the two (*Δ*ED50). This uncovered a strong, positive correlation between the predicted and observed *Δ*ED50 values, thus validating the predictive sequence features revealed by the fine-tuned ChromBPNet model (Figure 5C).

As an example, substantial buffering effects were observed with the *PTCHD4* enhancer, where converting three single E-boxes/low-affinity Coordinator instances to high-affinity Coordinators decreased ED50 from 33.9 to 19.8 and increased activity at 100% TWIST1 dosage by ∼14.5-fold (Figure 5D). In the opposite direction, converting two high-affinity Coordinator instances to a double E-box and low-affinity Coordinator in the buffered *MSR1* enhancer increased ED50 from 19.1 to 38 and decreased activity by ∼2.5 fold (Figure S7C). However, this correlation between buffering effects of mutations and increases in activity at 100% dosage was not the case across all mutants tested (Figure S7D). For example, insertion of 5 single E-box instances into the moderately buffered *IMPG1* enhancer increased ED50 from 18 to 45.3 with minimal change in activity at 100% TWIST1 dosage (Figure 5E). Together, these results indicate that sensitivity to TF dosage is not a simple correlate of unperturbed accessibility or enhancer activity, with a partially distinct sequence logic that can be experimentally modulated in a model-guided fashion.

### TF-nucleosome competition can explain the sensitizing effect of low-affinity sites

We sought to build biophysical intuition for why, as observed for TWIST1, low-affinity sites have a sensitizing effect when added in the vicinity of high-affinity, buffering sites. We considered a previously developed model^28^ in which competition between TFs and nucleosomes for binding to DNA (TF-nucleosome competition) induces TF cooperativity, referred to as ‘nucleosome-mediated cooperativity.’ TF-nucleosome competition through suppression of TF binding to nucleosome-bound DNA is controlled by a single parameter *c*, defined as the ratio of TF motif affinities between the nucleosome-free and -bound states. Given TF and nucleosome binding constants (the latter estimated from experimental data^28^), the model allows for derivation of steady state RE accessibility (inverse of nucleosome occupancy) as a function of TF concentration using a statistical mechanics approach.

We first implemented a simple case of this model (Figure 6A) with one high-affinity site, varied the K_D_ of a second site such that it went from high-affinity to low-affinity to non-existent (higher K_D_ meaning lower affinity), and calculated accessibility dosage response curves under varying degrees of TF-nucleosome competition by tuning *c*. We found that without TF-nucleosome competition (*c* = 0), the addition of any second site (high-or low-affinity) resulted in a more buffered dosage response (lower ED50) (Figure 6B), in contrast to the effects we observed from the ChromBPNet model and validated with enhancer reporter assays. In contrast, under strong competition (*c* = 0.01, i.e. TF binding has 100-fold lower affinity in the nucleosome-bound state), the addition of a second site with at least ∼10-fold higher K_D_ than the first site increases ED50 (Figure 6B), replicating the sensitizing effect we previously observed. With weak TF-nucleosome competition (*c* = 0.001), the second site had to be relatively weaker (∼1000-fold) in order to have a sensitizing effect, but results were qualitatively similar (Figure S8A).

**Figure 6.**
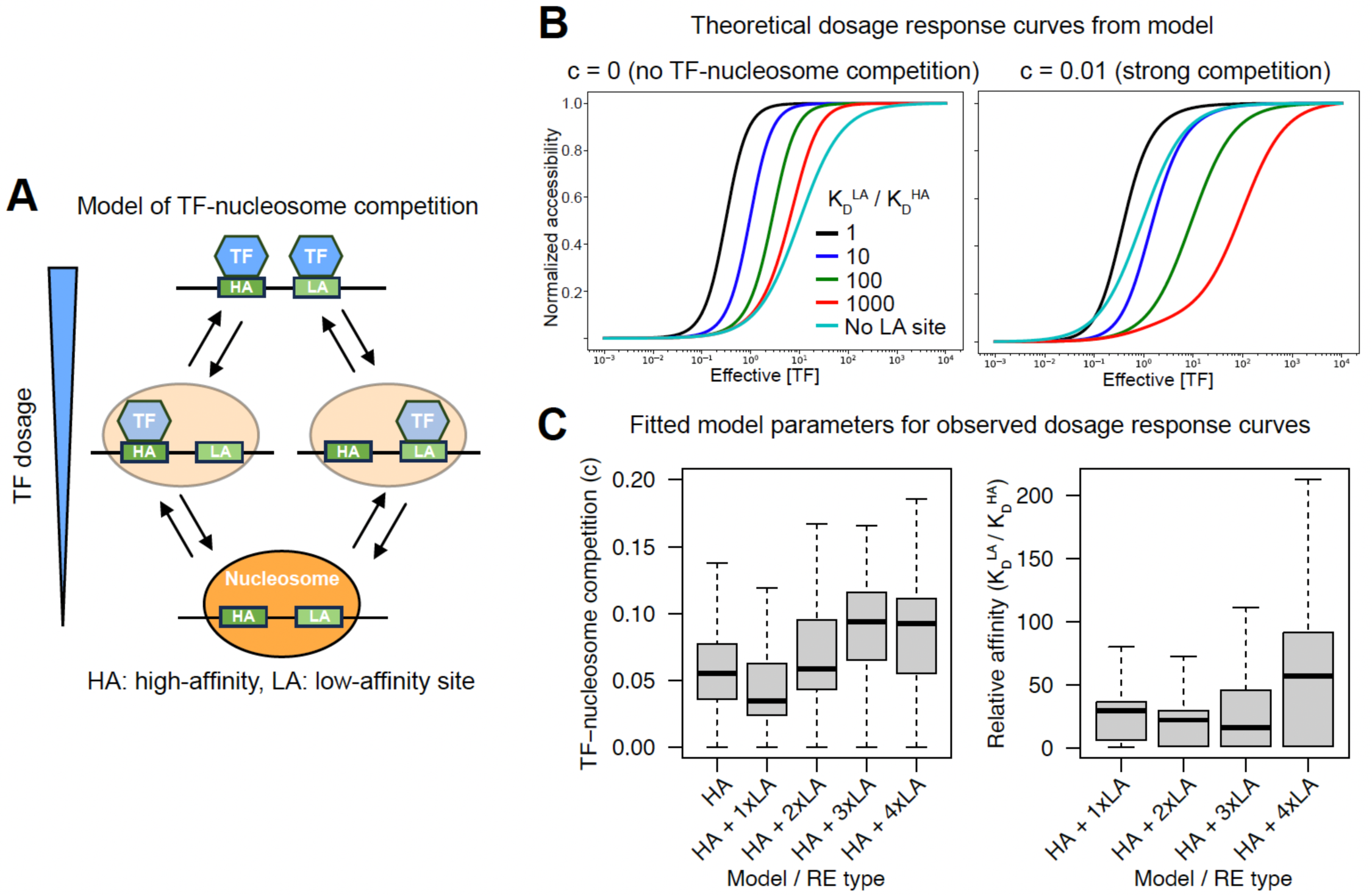
TF-nucleosome competition can explain the sensitizing effect of low-affinity sites. (A) Schematic of model of TF-nucleosome competition originally proposed by Mirny^28^. (B) Effect of low-affinity site (colors) or no site (grey) on theoretical dosage curves under model with no TF-nucleosome competition (left) or strong TF-nucleosome competition (right). The sensitizing effect of low-affinity sites (shift to right relative to grey curve) is only observed with competition. (C) Values of *c* (left) or low/high-affinity site K_D_ obtained by fitting model to observed dosage response curves for REs containing only high-affinity Coordinator sites (HA) or a mix of high-affinity sites and the indicated number of low-affinity Coordinator sites (HA + 1/2/3xLA). Numbers of points from left to right in left plot: 592, 925, 525, 360; from left to right in right plot: 925, 525, 360.

We next sought to fit this model of TF-nucleosome competition directly to our experimental dosage response curves to assess which values of *c* and relative sites affinities were best supported by real data. To simplify model fitting, we first fit to dosage response curves (parametrized by the ED50 and Hill coefficient) of 540 REs only containing high-affinity, buffering Coordinator motifs. We obtained the best fits with effective numbers of sites (n) ≥ 3 (Figure S8B), which yielded a median *c* of 0.022 (Figure 6C). We then held the effective number of high-affinity sites fixed at 3 and fit four types of models, each with one to three additional low-affinity sites, based on the number of observed low-affinity (sensitizing) sites in the RE (total 1,810 REs). The median *c* value of each model type ranged from 0.033 to 0.087, and the median K_D_ ratio between the low- and high-affinity sites ranged from ∼13-fold to ∼29-fold (Figure 6D). These fitted values support the scenario of strong TF-nucleosome competition analyzed theoretically above and are of the same order of magnitude as the ≥ 5-fold difference in affinity between canonical and degenerate Coordinator sequences suggested by our previous EMSA results^14^. We obtained similar fitted values when fixing the number of low-affinity sites in the model rather than matching to the number in each RE (Figure S8C). Together, these results suggest that TF-nucleosome competition can explain the sensitizing effect of low-affinity sites when added to REs containing high-affinity sites.

## Discussion

Here, we have used transfer learning to reveal the sequence features underlying the RE response to SOX9 or TWIST1 dosage. This approach can quantitatively predict both the magnitude and shape (buffered or sensitive) of the RE response. Interpreting the models revealed that, consistent with our previous work, composite or distinct motifs allowing for heterotypic TF binding lead to buffered responses. For TWIST1, our approach discovered low-affinity sequences that lead to sensitive responses even in the context of other buffering sequences. These sensitizing sequences are evolutionarily conserved, show the predicted effects when tested in enhancer reporter assays, and can be explained by TF-nucleosome competition.

Our study adds to the growing body of work demonstrating the importance of low-affinity TF binding sites for precise control of RE activity and gene expression. Low-affinity binding by TFs has been shown to confer specificity in both developmental^29,30^ and synthetic^31^ systems, and increases in affinity of such sites can result in ectopic expression and developmental phenotypes^32,33^. Our results broadly agree with these studies, as REs consisting solely of low-affinity TWIST1 binding sites have higher ED50s and require higher concentrations of TWIST1 to be active. Our observations further indicate that low-affinity sites play an important role in setting the shape of the TF dosage-response curve even in REs with other, high-affinity sites.

While our approach demonstrated the importance of low-affinity TWIST1 sites for TWIST1 dosage response, the models did not identify low-affinity (i.e. single) SOX9 sites as important for sensitivity to SOX9 dosage. This could be due to the fewer number of training examples for SOX9 sensitivity, but the fact that single SOX sites were not discovered as predictive of SOX9 full depletion effect, which uses much more training examples, while low-affinity Coordinators were discovered in the TWIST1 full depletion models, suggests that such data limitation is an unlikely explanation. Another possibility is that in CNCCs, single SOX motifs are of such low affinity that they effectively function as having no binding site at all. Consistent with this notion, previous *in vitro* binding assays showed highly cooperative SOX9 binding to the 3-5 bp palindrome, such that binding to one site enhanced binding to the second by >100-fold^25^. The only other motifs predictive of sensitivity to SOX9 dosage were those for the AP-1 TFs (dimers of the JUN, FOS, or ATF proteins). Previous studies have shown that SOX9 and AP-1 physically interact and bind overlapping genomic regions in chondrocytes^34^. It is possible that the AP-1 sites effectively function as low-affinity SOX9 sites by indirectly recruiting SOX9, but further experiments are needed to test this hypothesis.

Using a previously proposed model of TF-nucleosome competition, we showed that such competition is sufficient to explain the sensitizing effect of low-affinity binding sites. This model assumes a distance of up to ∼150-200 bp (the approximate DNA length of a single nucleosome) between low- and high-affinity binding sites, an assumption that holds true for a plurality (∼40%) of the low-affinitty sites studied here, with the majority of the remaining sites located within one or two nucleosomal lengths (Figure S8D). The 150-200 bp range of the TF-nucleosome competition model could be increased beyond a single nucleosome by inter-nucleosomal interactions^35^ or TF-mediated recruitment of chromatin modifiers. Indeed, a recent study of single-molecule TF and nucleosome occupancy added out-of-equilibrium kinetics (such as those induced by modifier recruitment) to explain observed TF co-binding as a function of nucleosome occupancy^36^. Explicitly modeling the effect of multi-nucleosome conformations and/or chromatin modifier recruitment on accessibility dosage response curves therefore represents an important future avenue.

Our results also provide a potential explanation for phenotypic specificity associated with TF dosage perturbations in human traits and diseases, despite widespread TF co-binding at REs. We observed that dosage-sensitive REs are largely non-overlapping between TWIST1 and SOX9, even among likely direct targets of both TFs. In this scenario, quantitative reductions in TWIST1 or SOX9 dosage preferentially impact largely distinct sets of REs, which in turn may regulate distinct genes and ultimately downstream phenotypes. In effect, these results extend our previously proposed model of phenotypic specificity at distinct SOX9 dosages^11^ to also explain how distinct phenotypes can arise from dosage perturbations of different TFs active in the same cell type.

Our study highlights the power of applying deep learning to epigenomic data gathered from perturbed states as a means of learning important cis-regulatory logic that would not be apparent in steady-state data. Many of the low-affinity, sensitizing sequences that our approach discovered would not be detected by traditional motif matching approaches or deep learning models of unperturbed chromatin accessibility. This additional layer of the cis-regulatory code, one that sets the response to TF dosage changes, could be modulated by trait-associated sequence variants discovered in genome-wide association studies (GWAS). The types of models we have described here could therefore ultimately aid in interpreting and fine-mapping GWAS variants, particularly those that cannot readily be explained by effects on chromatin state in an unperturbed setting.

## Methods

### Cell culture

Female H9 (WA09; RRID: CVCL_9773) hESCs were obtained from WiCell and cultured in either mTeSR1 (Stem Cell Technologies 85850) for at least one passage before differentiation into CNCCs or mTeSR Plus (Stem Cell Technologies 100–0276) for gene editing, single-cell cloning, expansion and maintenance. hESCs were grown on Matrigel growth factor reduced basement membrane matrix (Corning 354230) at 37 °C. hESCs were fed every day for mTeSR1 or every 2 days for mTeSR Plus and passaged every 5–6 days using ReLeSR (Stem Cell Technologies 05872).

### Differentiation of hESCs to CNCCs

hESCs were grown for 5–6 days until large colonies formed, and then they were disaggregated using collagenase IV and gentle pipetting. Clumps of about 200 hESCs were washed in PBS and transferred to a 10 cm Petri dish in neural crest differentiation medium (1:1 ratio of DMEM-F12 and Neurobasal, 0.5× Gem21 NeuroPlex supplement with vitamin A (Gemini, 400-160), 0.5× N2 NeuroPlex supplement (Gemini, 400-163), 1× antibiotic–antimycotic, 0.5× Glutamax, 20 ng ml^−1^ bFGF (PeproTech, 100-18B), 20 ng ml^−1^ EGF (PeproTech, AF-100-15) and 5 μg ml^−1^ bovine insulin (Gemini Bio-Products, 700-112P)). After 7–8 days, neural crest emerged from neural spheres attached to the Petri dish, and after 11 days, neural crest cells were passaged onto fibronectin-coated 6-well plates (about 1 million cells per well) using Accutase (Sigma-Aldrich A6964) and fed with neural crest maintenance medium (1:1 ratio of DMEM-F12 and neurobasal, 0.5× Gem21 NeuroPlex supplement with vitamin A (Gemini, 400-160), 0.5× N2 NeuroPlex supplement (Gemini, 400-163), 1× antibiotic– antimycotic, 0.5× Glutamax, 20 ng ml^−1^ bFGF, 20 ng ml^−1^ bFGF EGF and 1 mg ml^−1^ BSA (Gemini)). After 2–3 days, neural crest cells were plated at about 1 million cells per well of a 6-well plate, and the following day cells were fed with neural crest long-term medium (neural crest maintenance medium + 50 pg ml^−1^ BMP2 (PeproTech, 120-02) + 3 μM CHIR-99021 (Selleck Chemicals, S2924; BCh medium)). After transition to BCh medium, CNCCs at subsequent passages were plated at about 800,000 cells per well of a 6-well plate. CNCCs were then passaged twice to passage 4, at which depletion experiments were carried out. For depletion experiments, dTAG^V^-1 (Tocris, 6914) at a range of concentrations was added to BCh medium, with an equivalent amount of dimethylsulfoxide (DMSO) as vehicle control.

### Flow cytometry

CNCCs were collected for intracellular staining and flow cytometry using Accutase treatment for 5 min at 37C then washed twice in PBS. Cells were fixed in 4% Paraformaldehyde in PBS for 10 min at RT, washed twice with PBS, and then permeabilized in 0.1% TritonX 100 in PBS for 10 min at RT. Cells were then resuspended in blocking buffer (1% BSA, 3% donkey serum in PBS) and blocked for 40 min at RT, flicking tubes every 10 min to resuspend settled cells. Fixed, permeabilized, and blocked cells were incubated with primary antibody (V5) diluted 1:100 in blocking buffer and incubated on ice for 60 min with flicking every 10 min, followed by two PBS washes and staining with secondary antibody for 60 min at RT. Cells were finally washed two additional times with PBS prior to flow cytometry analysis, which used to measure V5 staining intensity and/or mNeonGreen fluorescence (for *SOX9*-tagged CNCCs) after excluding doublets and debris based on forward and side scatter (Beckman Coulter Cytoflex). Fluorescence values were summarized per biological replicate using geometric means.

### ATAC-seq collection and library preparation

CNCCs were incubated with BCh medium containing 200 U ml DNase I (Worthington, LS002007) for 30 min and collected using Accutase. Viable cells were counted using a Countess Automated Cell Counter (Invitrogen), and 50,000 viable cells were pelleted at 500 RCF for 5 min at 4 °C and resuspended in ATAC-resuspension buffer (10 mM Tris-HCl pH 7.4, 10 mM NaCl, 3 mM MgCl_2_ in sterile water) containing 0.1% NP-40, 0.1% Tween20 and 0.01% digitonin and incubated on ice for 3 min. Following wash-out with cold ATAC-resuspension buffer containing 0.1% Tween20, cells were pelleted and resuspended in 50 μl transposition mix (25 μl 2× TD buffer, 2.5 μl transposase (100 nM final), 16.5 μl PBS, 0.5 μl 1% digitonin, 0.5 μl 10% Tween20, 5 μl H_2_O) and incubated for 30 min at 37 °C with shaking. The reaction was purified using the Zymo DNA Clean & Concentrator kit and PCR-amplified with NEBNext High-Fidelity 2× PCR Master Mix (NEB, M0541L) and primers as defined in Corces et al. Libraries were purified by two rounds of double-sided size selection with AMPure XP beads (Beckman Coulter, A63881), with the initial round of 0.5× sample volume of beads followed by a second round with 1.3× initial volume of beads. Library size distributions were confirmed by separation on a PAGE gel and staining with SYBRGold and pooled on the basis of quantifications from Qubit dsDNA High Sensitivity Kit. Pooled libraries were sequenced using the Novaseq 6000 platform (2 × 150 bp).

### Luciferase assays

CNCCs were transfected with the appropriate plasmids immediately following passaging to passage 5 in 48-well plates. For *TWIST1-*tagged CNCCs, dTAG^V^-1 treatment to titrate TWIST1 dosage were started at the time of transfection, whereas for *SOX9-*tagged CNCCs, dTAGV-1 treatment was started 24h prior to passaging to p5 and transfection. Four independent transfections were performed for each dTAG^V^-1 concentration, with each well receiving 5ng of pGL3 plasmid, 0.25ng of control pRL firefly renilla plasmid, 44.25 μL carrier DNA (circularized pUC19 plasmid) and 0.3 μL Fugene 6 in 25 μL of optimum. The pGL3 plasmid contains the firefly luciferase gene driven by an SV40 promoter with either a control SV40 enhancer downstream, or a test enhancer sequence cloned upstream (Promega), the pRL plasmid acts as a transfection control with Renilla luciferase driven by an upstream CMV enhancer and CMV promoter (Promega). Test enhancers were cloned by either PCR of genomic DNA with primers containing NheI and XhoI restriction sites, or synthesized by Twist Biosciences with NheI and XhoI flanking restriction sites, and ligated into NheI and XhoI-digested pGL3 vector. 24 hours after transfection, cells were washed in PBS, and lysed in 65 μL 1X passive lysis buffer (in PBS) for 15 min (Promega). 20 μL lysate was then transferred to an opaque flat-bottomed plate for reading with a luminometer (Veritas). An automated injector added 100 μL LARII reagent and the well was read using the following parameters: 2 s delay, 10 s integration. 100 μL Stop-and-Glow reagent was then injected into the well and read using the same parameters. Empty vector and the EC1.45 min1-2 enhancer (shown to be strongly active in CNCCs by Long et al) were included in each experiment as negative and positive controls, respectively.

## Quantification and statistical analysis

### ATAC-seq preprocessing

Reads were trimmed of Nextera adapter sequences and low-quality bases (-Q 10) using skewer v0.2.2 and then mapped to the hg38 analysis set (human) using Bowtie2 v2.4.1 with the options --very-sensitive -X 2000. Reads were deduplicated with <u>samtools</u> v1.10 markdup and uniquely mapped reads (-q 20) mapped to the main chromosomes (excluding mitochondria and unplaced contigs) were retained using samtools view. Read ends were shifted inward 5 bp (+5 bp on + strand, -5bp on – strand) for each fragment, and then counts of reads in each sample overlapping the reproducible peak set of 151,457 REs from Naqvi et al were generated using bedtools.

### Modeling of TF dose-response curves

TWIST1-dependent REs were defined by differential accessibility between undepleted and fully depleted TWIST1 concentrations was carried out using DESeq2 v1.32.0, with CNCC differentiation batch as a covariate and raw counts as input. SOX9-dependent REs from Naqvi et al were used. RE ATAC CPM values were first TMM-normalized using the edgeR package v3.34.0. For each TWIST1/SOX9-dependent RE, were corrected for differentiation batch effect by linear regression using the lm() function. Differentiation-corrected CPM values were scaled by dividing by the maximum absolute value across samples. Sample outliers, defined as *z*-score greater than 3, were removed from the analysis of that RE/gene. The data were then to the Hill equation using the drm() function in the drc R package v3.0-1. A two-parameter Hill equation (that is, with minimum and maximum fixed as the mean CPM at full or no depletion, respectively) was used unless a three-parameter Hill equation with fixed minimum but free maximum was a better fit (decrease in AIC > 2 relative to the two-parameter model); for these genes/REs, the three-parameter Hill was used. The Hill exponent as fitted was extracted, but for ED50 we used a modified calculation, calculating the TF dosage at which the normalized ATAC signal reached 50% of the signal at 100% TF dosage (rather than the theoretical maximum). This essentially caps the ED50 at 100% and avoids instability in high ED50 estimates especially when the three-parameter Hill was used.

### Prediction of RE responsiveness from DNA sequence

#### Definition of training, testing, and validation sets

For predicting effect of full TF depletion, all 151,457 RE peaks were used and divided into training, test, and validation sets based on the chromosome-level “fold 0” training, testing, and validation split from the ChromBPNet package. For predicting ED50, the same fold split was used but only among the likely direct target REs of each TF. For SOX9 this was the “Rapid down” class of REs from Naqvi et al (i.e. downregulated in accessibility after 3h full SOX9 depletion), and for TWIST1 this was defined as REs bound by TWIST1 ChIP-seq and downregulated at 24h.

#### Baseline approaches

For baseline approaches to predicting effect of full TF depletion or ED50, we encoded sequence information by quantifying known PWM matches (PWMs as defined in the HOCOMOCO v11 core set) in a 200 bp window around each RE ATAC peak summit. PWMs were obtained from the HOCOMOCO v11 core PWM set, and matched to sequences using MOODS v1.9.4.1 with -p 0.01 as a permissive cutoff. For each RE, the best PWM match (as determined by the highest MOODS match score) was stored and quantified with the MOODS match score. PWMs with no reported match were set to 0. GC and CpG content as well as unperturbed ATAC-seq signal (quantified as log_10_(baseMean) output from the DESeq model) were added as additional predictors. The matrix of predictors for the training and test sets were separately centered and scaled to have mean 0 and standard deviation 1. LASSO regression was performed using the cv.glmnet package in r with alpha 0.01 and nlambda 50. Random forest regression was performed with the randomForest package in r with ntree 100.

#### Deep learning model pretraining and fine-tuning

The pretrained deep learning model was obtained by running the full ChromBPNet v0.1.1 pipeline with default parameters on a consolidated BAM file of all unperturbed TF ATAC-seq samples. The 151,457 RE peak set from Naqvi et al was used and a corresponding background peak set was created using chrombpnet prep nonpeaks. The “fold 0” training, testing, and validation split was used. Next, the model was fine-tuned with the effect size of full TF depletion or ED50 from the relevant training set. Learning rate was set to 1e-3 as in the original ChromBPNet training, with training for 10 epochs. The best-performing model (lowest loss on the validation set) was used. The same loss functions as the pretrained model were used, except the weight for the multinomial NLL loss (for the base-resolution profiles) was set to 0. Reverse-complemented sequences were used as data augmentation.

#### Model interpretation and motif matching

Contribution scores for both the pretrained and fine-tuned ChromBPNet models were extracted using chrombpnet contribs_bw with -pc counts, and TF-MoDISCO was run for motif discovery using the chrombpnet modisco_motifs command with -N 1000000. Top CWMs output from TF-MoDISCO were matched to their genomic locations by adapting a previously described procedure^6,7^ that considers both the Jaccardian similarity between a CWM and a test sequence as well as that sequence’s overall contribution score. As in Brennan et al, because we were interested in CWM matches corresponding to low-affinity motifs, mapping thresholds were lowered to mapping the motif if the CWM Jaccard similarity percentile was equal to or greater than 10% and if the total absolute contribution percentile was equal to or greater than 0.5%. After mapping, motifs were filtered for redundant assignment of palindromic sequences and overlapping peaks; if multiple different CWMs matched the same sequence (as was frequent with partial and degenerate Coordinators), the CWM with the highest Jaccard similarity score (multiplied by CWM length to account for the fact that higher match scores are more likely with short motifs) was chosen.

#### Analyses of evolutionary constraint

Basepair-level PhastCons and phyloP scores from alignment of 30 primate genomes were obtained from https://hgdownload.soe.ucsc.edu/goldenPath/hg38/phastCons30way/ and https://hgdownload.soe.ucsc.edu/goldenPath/hg38/phyloP30way/ respectively, and averaged over buffering or sensitizing CWM occurrences as indicated. The shuffled CWM occurrence set was generated with bedtools shuffle on a sufficiently large number of arbitrary regions within the set of TWIST1/SOX9-dependent REs, and then subtracting the true CWM occurrences. Selection estimates based on both within-species polymorphism and between-species divergence were obtained from the INSIGHT^26^ web tool (http://compgen.cshl.edu/INSIGHT/).

#### Modeling of chromatin accessibility

The nucleosome-mediated cooperativity (NMC) model proposed by Mirny, 2010 considers a DNA region that is either in a nucleosome-bound or open state. Assuming that there are n number of transcription factor binding sites in this region, the transcription factor can either bind in the nucleosome-bound or open state. However, there is a suppression of transcription factor binding in the nucleosome-bound state due to the energy cost of DNA unwrapping (i.e. TF-nucleosome competition). In the simplest form, there are three dimensionless parameters to describe the system: the equilibrium between nucleosome-bound and open state in the absence of transcription factors denoted as L, TF-nucleosome competition denoted as c, and effective protein concentration denoted as *α*. Using this model, the nucleosome occupancy (Y_N_) can be assayed as a function of protein concentration to assess chromatin accessibility. To study the effect of combining high-affinity and low-affinity transcription factor binding sites on accessibility, we assumed a region of DNA with a high- and a low-affinity sites. Accessibility is then calculated using the following equation:

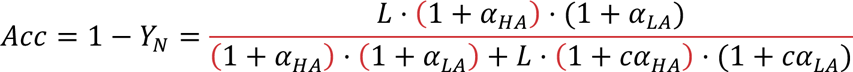

Where *α*_LA_ and *α*_HA_ are the effective protein concentration for the strong and weak sites, defined as 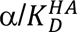 and 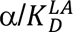. The model parameters were set to L=10^3^, c=0 or 0.01 or 0.1, and *α* was titrated between 10^-5^ to 10^5^ to calculate the ED50 of the dosage response curves. To fit observed accessibility dosage response curves to the above model, python’s SciPy curve_fit library was used. First, the ED50 and Hill coefficient from the reporters containing only high-affinity sites was used to generate response curves that were fitted to the following equation:

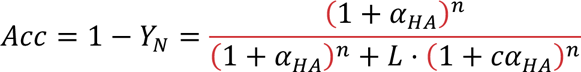

L was fixed at 10^3^. The mean squared error of the fit was calculated for n of 1 to 7 and it was determined that optimal fitting is achieved for n > 2, and thus we chose n^HA^ = 3. Next, the REs containing high- and low-affinity sites was fit to the following equation:

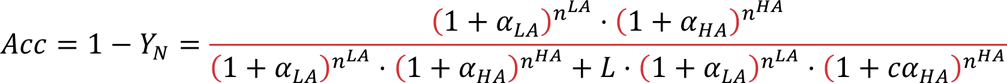

Where n^LA^ refers to the modeled number of low-affinity sites and is either matched to the number of sensitizing elements present in that RE, or is fixed at different values for all REs. Poor model fits (mean squared error > 0.001 or negative fitted values of c) were removed from further analysis.

## Data availability

The raw sequencing files generated during this study are available on the Gene Expression Omnibus (accession number GSE267008); corresponding processed data and trained deep learning models are available on Zenodo^37^. TF-binding motifs were obtained from HOCOMOCO v11 (https://hocomoco11.autosome.org/). Plasmids generated in this study will be deposited in Addgene. All other reagents are available upon request to S.N.

## Code availability

The ChromBPNet package is available on GitHub (https://github.com/kundajelab/chrombpnet). Additional code used for sequencing data analysis and finetuning of ChromBPNet models, as well as output of pretrained and fine-tuned ChromBPNet models, is available on Zenodo^37^.

## Acknowledgements

We thank Surag Nair, Jordan Valgardson, and Tony Zeng for advice on the transfer learning approach. This work was supported by a Helen Hay Whitney Fellowship and NIH grant K99 DE032729 to S.N.; an HHMI-Damon Runyon Cancer Research Foundation fellowship (DRG-2420-21) to S.K.; the Stanford Graduate Fellowship and the NSF Graduate Research Fellowship (DGE-2146755) to S.T; NIH grant R01 HG008140 to J.K.P; and the NIH grant R35 GM131757, the Nomis Foundation, funding from the Howard Hughes Medical Institute, a Lorry Lokey endowed professorship, and a Stinehart Reed award to J.W.

---

**Figure S1.**
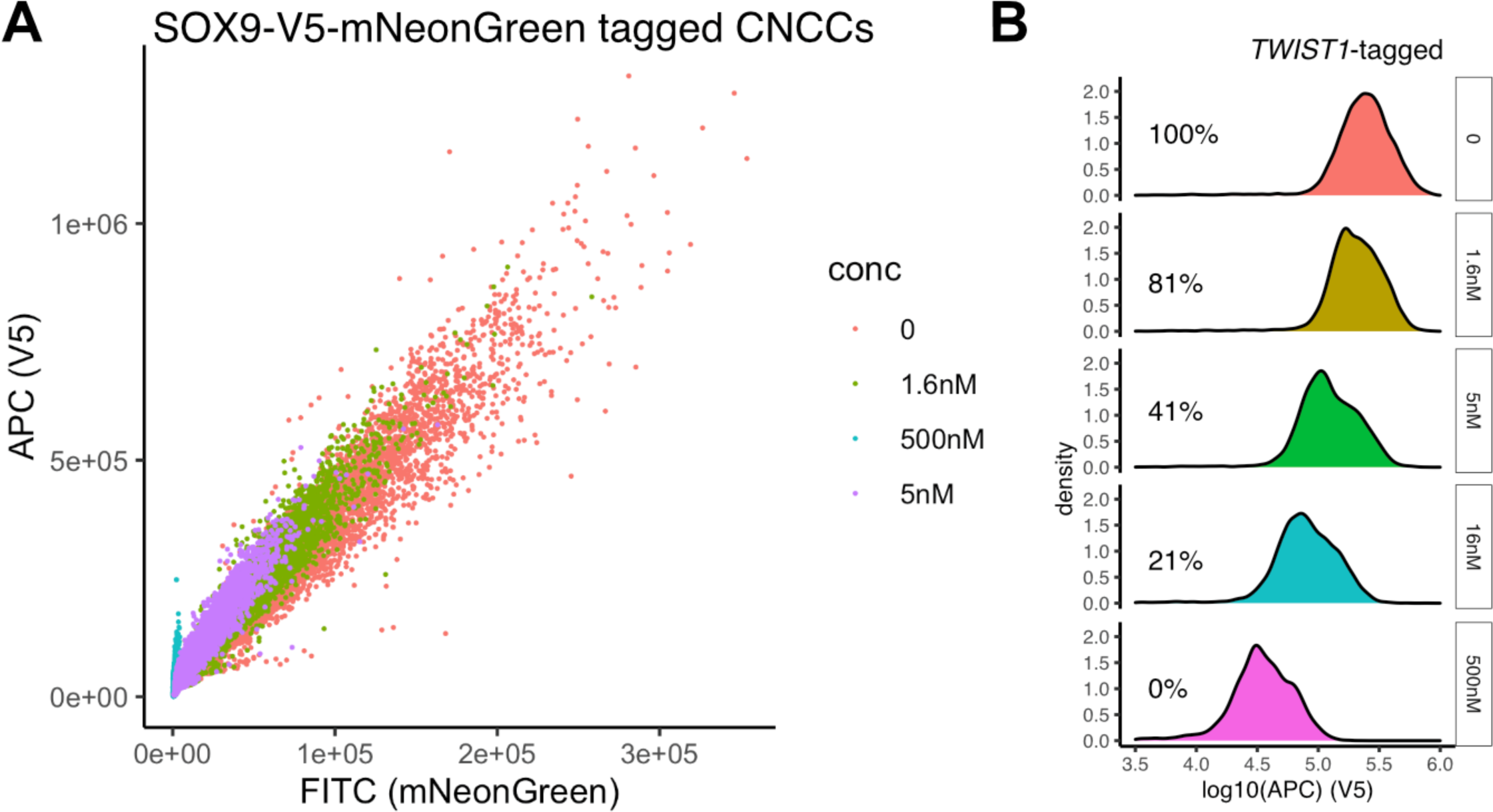
Precise modulation of TWIST1 dosage. (A) Comparison of V5 (y-axis) and mNeonGreen (x-axis) signal in single SOX9-tagged cells treated with different dTAGV-1 concentrations. (B) Second independent replicate of TWIST1 dosage modulation, as in Figure 1B.

**Figure S2.**
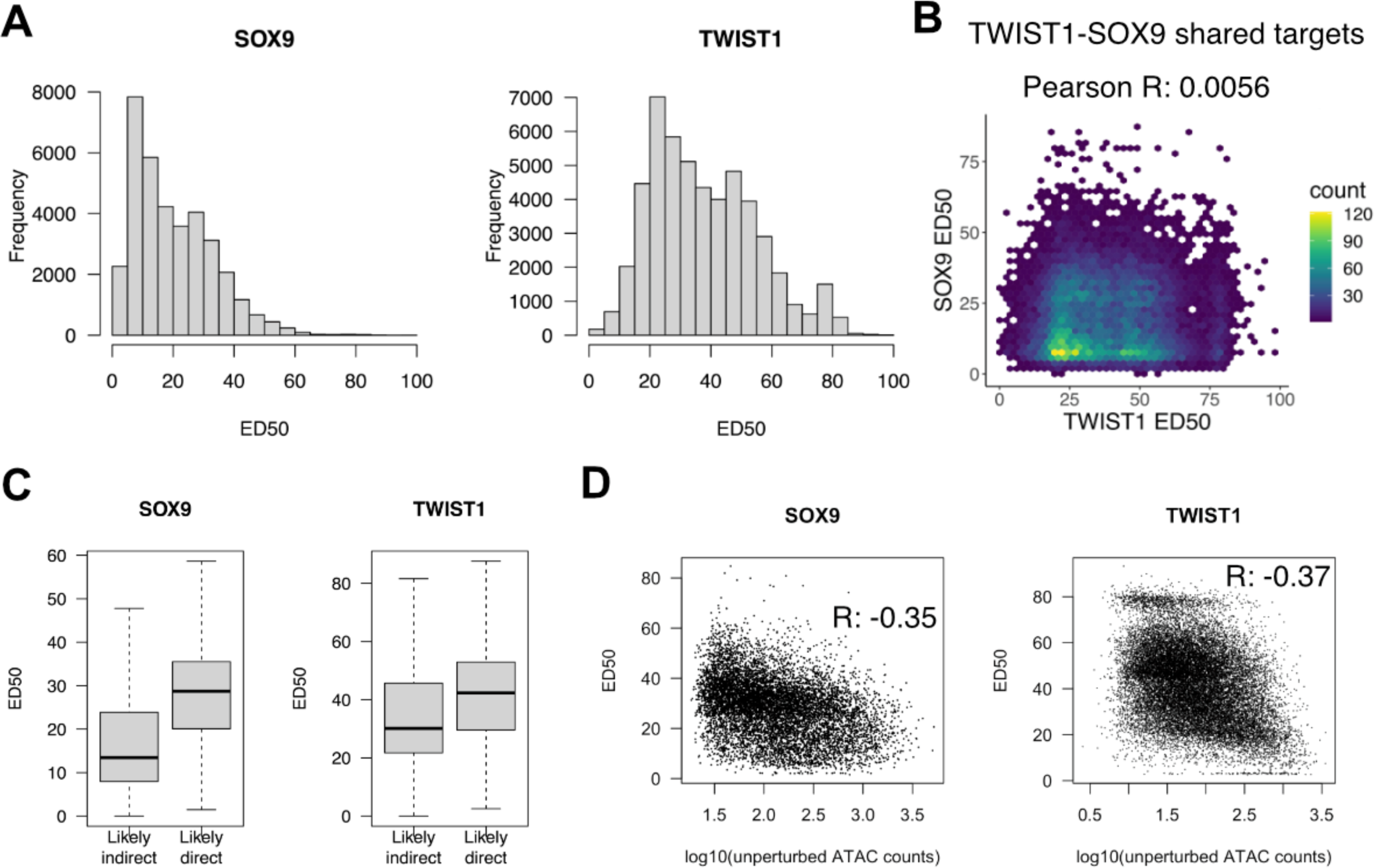
RE sensitivity to SOX9 and TWIST1 dosage. (A) Distribution of ED50 values among all SOX9 or TWIST1-dependent REs. (B) ED50 with respect to SOX9 dosage (y-axis) and TWIST1 dosage (x-axis) for all REs that are both SOX9- and TWIST1-dependent. (C) ED50 of likely direct or indirect SOX9 or TWIST1 targets. For SOX9, likely direct targets were defined as the 3h downregulated class as in Naqvi et al 2023, and for TWIST1, direct targets were defined as downregulated and containing a TWIST1 ChIP-seq peak. (D) Unperturbed accessibility (x-axis) versus ED50 (y-axis) for all likely direct SOX9 or TWIST1 targets.

**Figure S3.**
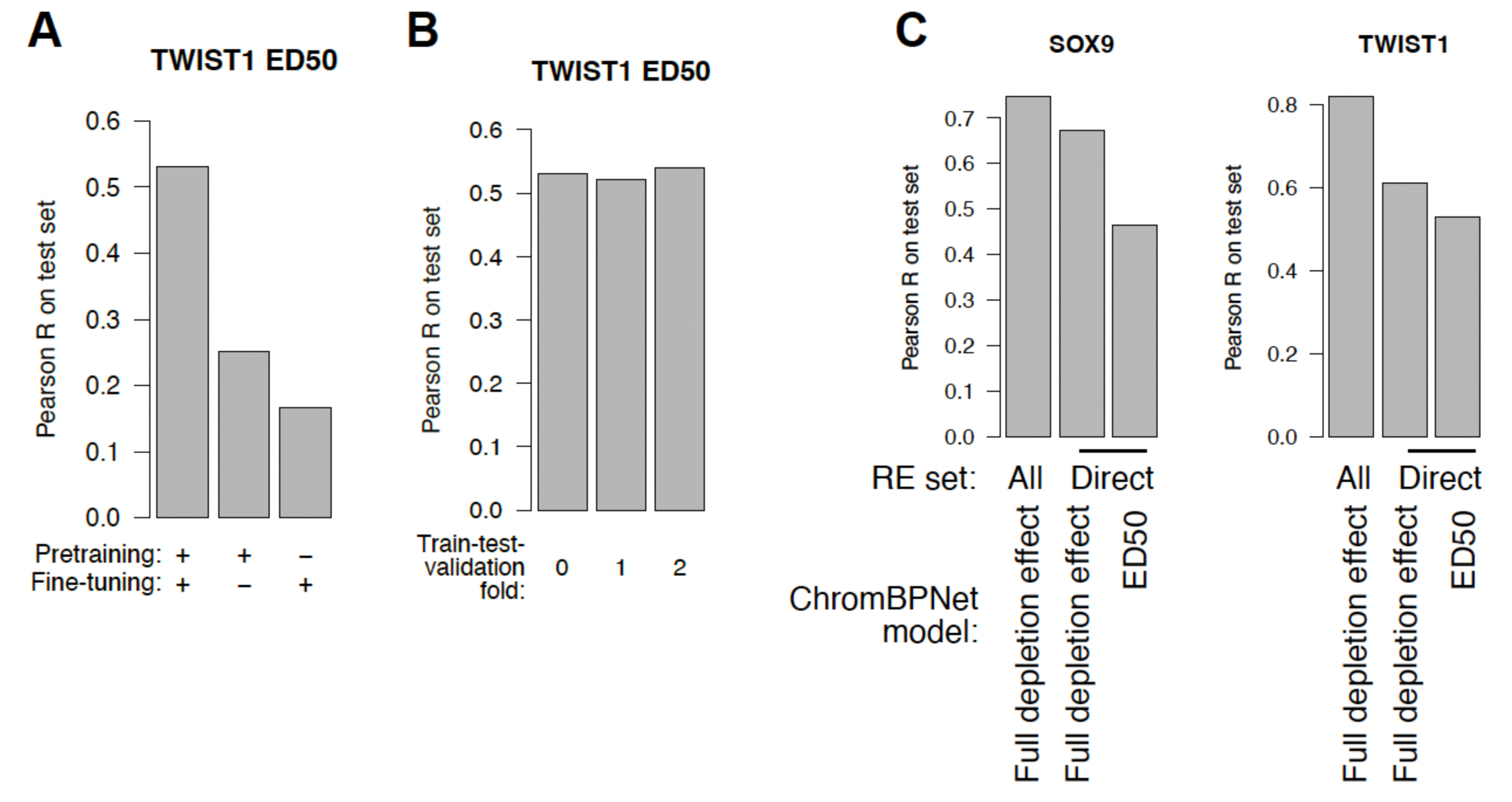
Prediction of effect of full TF depletion and ED50 from DNA sequence. (A) Prediction accuracy of TWIST ED50 prediction with and without pretraining or fine-tuning of ChromBPNet model. (B) Performance of pretrained and fine-tuned ChromBPNet model for predicting TWIST1 ED50 across three independent train-test-validation splits. (C) Decreased performance of ChromBPNet model for prediction effect of full SOX9 (left) or TWIST1 (right) depletion when predictions are subsetted to only direct RE targets (middle bar in each plot).

**Figure S4.**
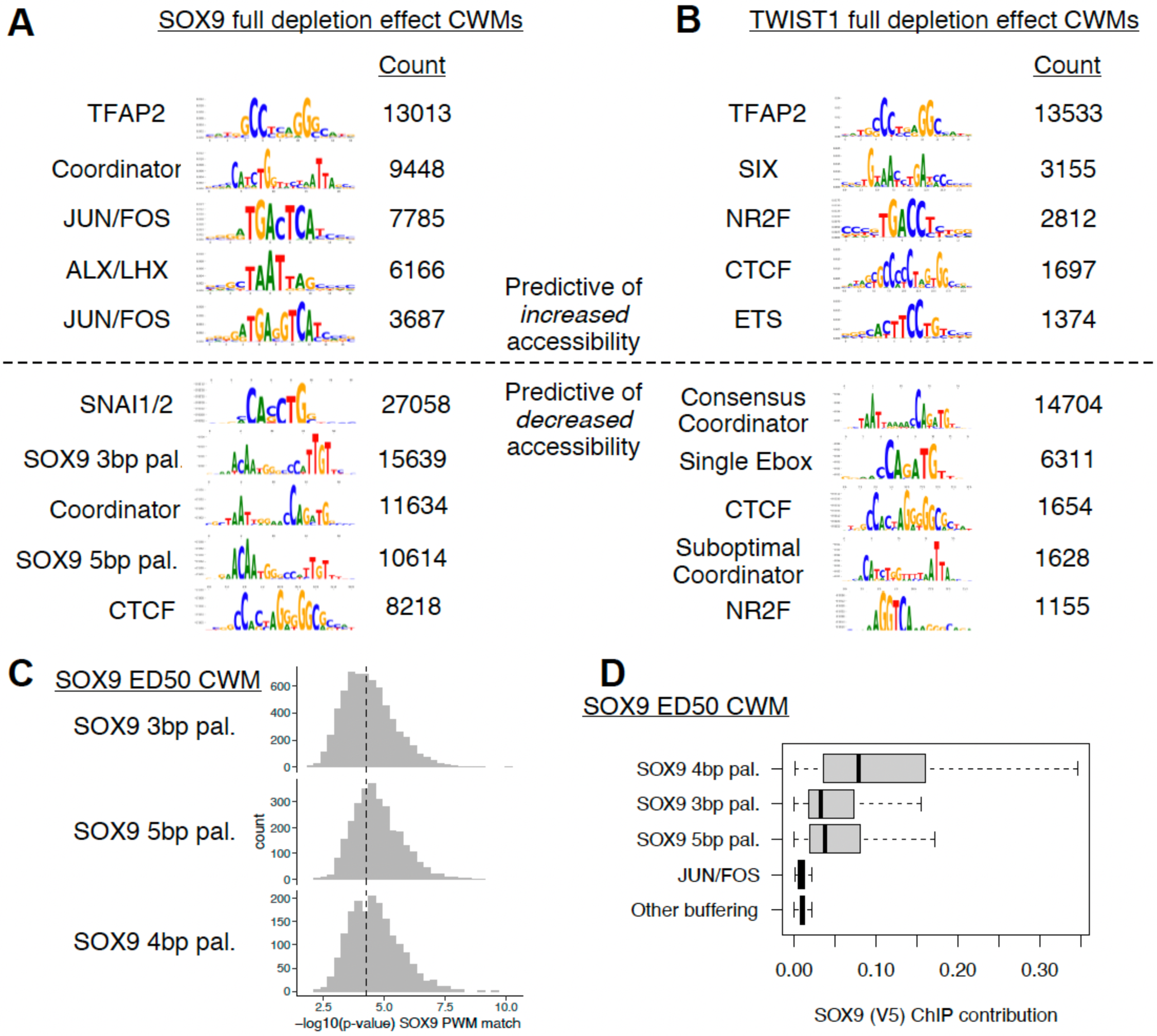
Sequence features predictive of the effect of full TF depletion on RE accessibility. (A,B) Top contribution weight matrices (CWMs) predictive of effect of full depletion of (A) SOX9 or (B) TWIST1 on RE accessibility. Number of individual occurrences of each CWM is indicated under the “count” column. (C) For all individual instances of the indicated CWMs predictive of SOX9 ED50 (rows), the strength of that sequence match to SOX9 palindrome position weight matrix (PWM) is shown (x-axis). (D) For the indicated CWMs, the distribution of contribution to SOX9 binding, estimated from BPNet on SOX9-V5 ChIP-seq, is shown.

**Figure S5.**
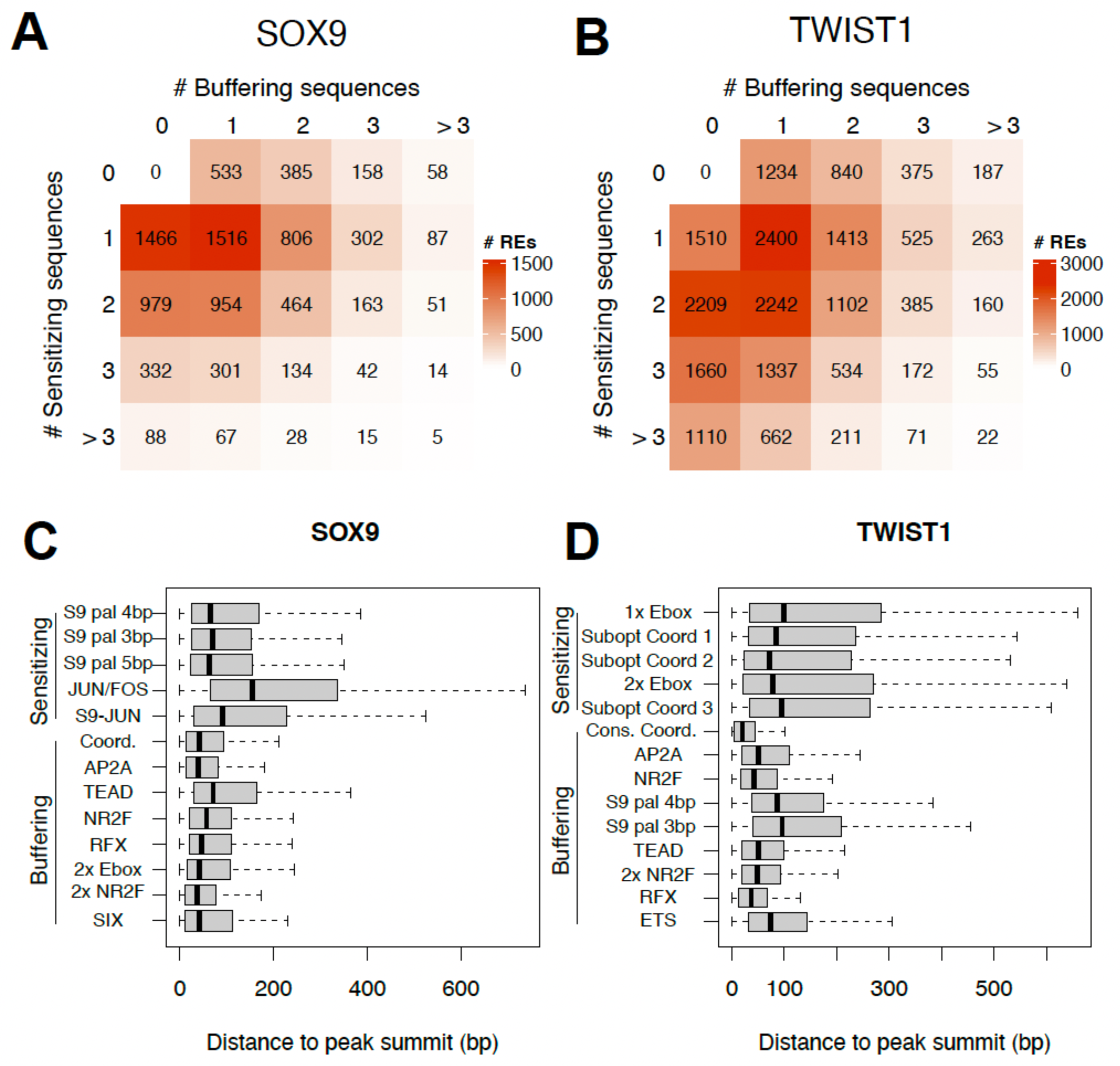
Additional features of buffering and sensitizing sequences. (A,B) The number of (A) SOX9 or (B) TWIST1 target REs with the indicated number of buffering (x-axis) or sensitizing (y-axis) CWM occurrences. (C,D), Distance to ATAC peak summit of individual types of sensitizing or buffering CWM occurrences for SOX9 (C) or TWIST1 (D).

**Figure S6.**
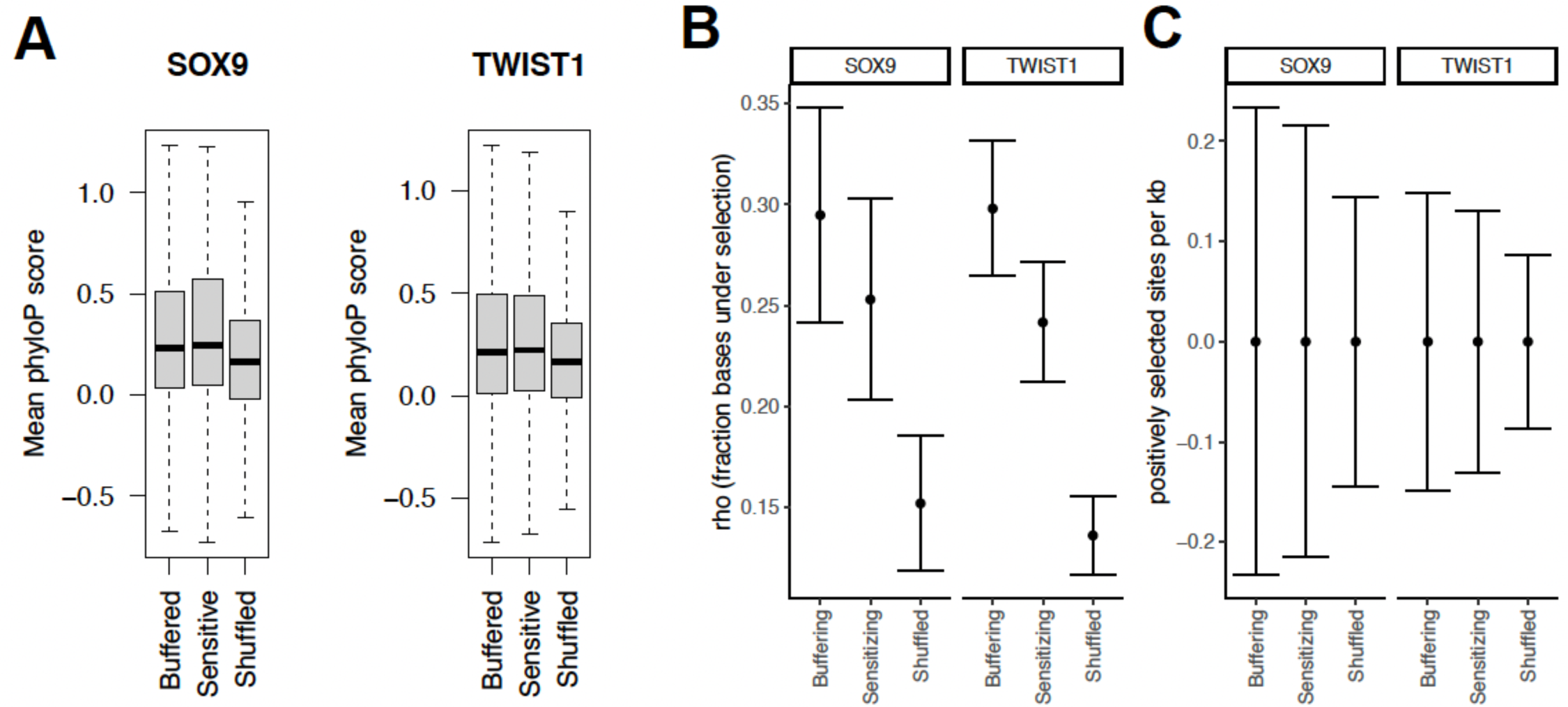
Signatures of selection at sensitizing and buffering sequences. (A) Mean phyloP score of buffering, sensitizing, or shuffled motif occurences for SOX9 or TWIST1 ED50. Positive phyloP scores mean more likely to be conserved, negative means more likely to be under positive selection. (B,C) Fraction of sites under weak negative selection (B) or frequency of sites under positive selection (C) for the same classes of motifs as in (A), estimated by INSIGHT.

**Figure S7.**
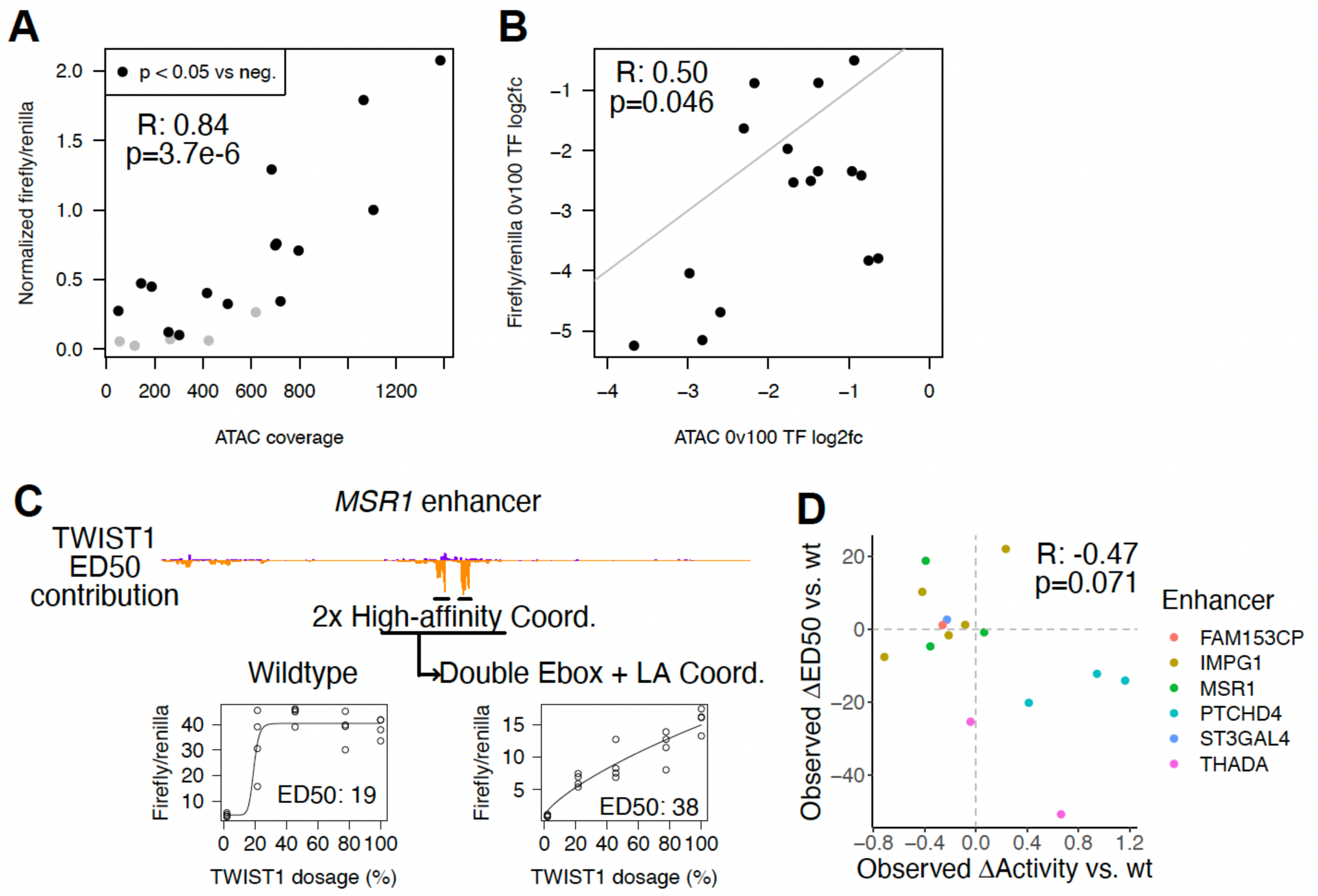
Dosage responses of wildtype and mutant REs in enhancer reporter assays. (A) Comparison of endogenous unperturbed accessibility (x-axis) and enhancer reporter activity (y-axis, normalized to positive control in each experiment) across 19 TWIST1- and SOX9-dependent REs (points). (B) For the REs in (A) with significantly higher activity than the negative control, the effect of full TF depletion on endogenous accessibility (x-axis) is compared to the effect of full TF depletion on enhancer reporter activity (y-axis). (C) Example of *MSR1* enhancer, where converting two high-affinity, buffering Coordinator motifs into a double E-box and low-affinity (LA) Coordinator motif has a sensitizing effect. (D) Comparison of changes relative to wildtype in reporter activity at 100% TF dosage (x-axis) and ED50 (y-axis) for tested mutant enhancers.

**Figure S8.**
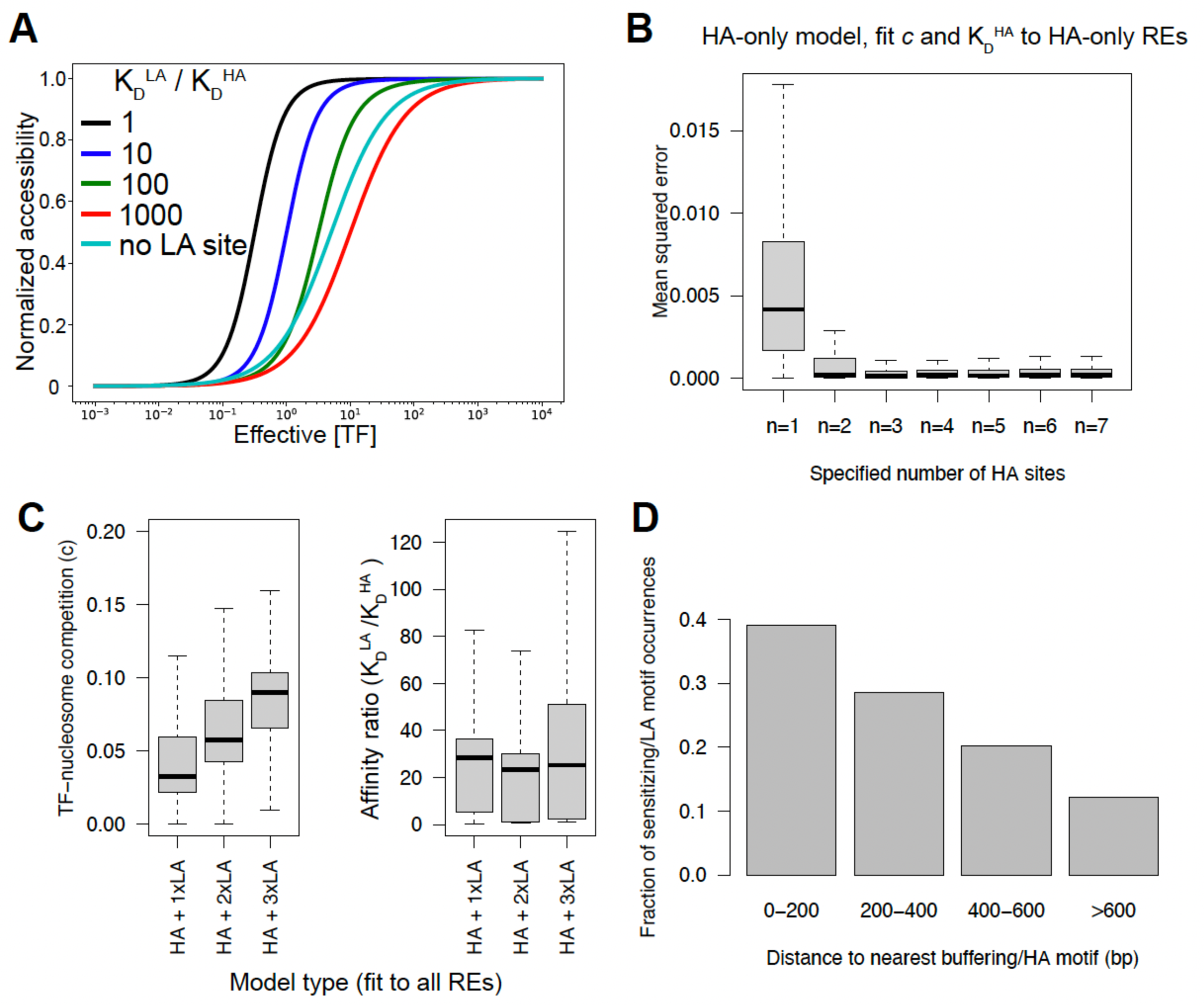
Theoretical and fitted instances of TF-nucleosome competition model. (A) Effect of low-affinity site (colors) or no site (grey) on theoretical dosage curves under model weak TF-nucleosome competition (*c* = 0.001). (B) Mean squared error of high-affinity (HA)-only model with specified effective number of HA sites (x-axis), fit to 1,291 REs. (C) Values of *c* or high/low-affinity site K_D_ obtained by fitting model to observed dosage response curves for REs a mix of high-and low-affinity Coordinator sites (HA + LA). All REs were fit with the indicated model rather than models matched to the number of LA REs in each. (D) For REs containing a mix of buffering, high-affinity (HA) and sensitizing, low-affinity (LA) motifs, the fraction of LA motifs within the indicated distance to the nearest HA motif is shown.

